# The fungal protein Jps1 facilitates unconventional protein secretion through a direct phosphoinositide interaction

**DOI:** 10.1101/2024.02.29.582524

**Authors:** Sanchi Dali, Michèle Schultz, Marian Köster, Michael Kamel, Max Busch, Wieland Steinchen, Sebastian Hänsch, Athanasios Papadopoulos, Jens Reiners, Sander H. J. Smits, Alexej Kedrov, Florian Altegoer, Kerstin Schipper

## Abstract

Protein secretion is indispensable for essential cellular processes in eukaryotic cells, contributing significantly to nutrient acquisition, defense or communication. Alternative pathways bypassing the endomembrane system collectively referred to as unconventional secretion are gaining increasing attention. A number of important molecules such as cytokines, fibroblast growth factor or viral proteins are being exported through these mechanistically diverse pathways. In the fungal model *Ustilago maydis*, cytokinesis-dependent unconventional secretion mediates export of the chitinase Cts1 via the fragmentation zone. This membrane-rich compartment is formed during cytokinesis between mother and daughter cells. Recently, we identified Jps1, a previously uncharacterized protein, as a crucial factor for Cts1 localization and export. Combining biochemical experiments and *in vivo* studies, we here uncover two pivotal features of Jps1: dimerization and phosphoinositide (PIP) binding. Our findings reveal that a conserved structural core domain mediates homodimerization, while surrounding flexible variable regions suggest potential diversification in different basidiomycete species. Jps1 does not harbor a canonical PIP-binding domain but instead specificity of the interaction with the preferred PIP PI(4,5)P_2_ is determined by basic residues. Importantly, loss of PI(4,5)P_2_ binding specificity results in mis-localisation, morphological defects and reduced extracellular Cts1 activity, particularly at low cell densities. This discovery sheds light on previously unknown key features of Jps1, elucidating its role in supporting Cts1 secretion, and representing a crucial step towards understanding the broader implications of unconventional secretion in eukaryotic cells.

## Introduction

Cells rely on an intricate selection of protein secretion pathways to fulfill vital functions essential for survival. These pathways are crucial for tasks like acquiring nutrients, building defense responses, and communicating with other cells. In eukaryotic cells, secretion was long thought to predominantly occur via the endomembrane system (1). Here, proteins destined for secretion are targeted to the Endoplasmic reticulum (ER) via N-terminal signal peptides (2–4). After entry of the ER through the Sec61 translocon, the proteins undergo folding and are eventually modified, for example by *N*-glycosylation and disulfide bonds (5). Besides this well-established secretion via the endomembrane system, various unconventional secretion routes where proteins are exported despite the lack of N-terminal signal sequences were unveiled in recent years (6–8). Unconventional secretion of soluble proteins can be grouped into mechanisms of transfer through the plasma membrane via direct translocation (type I) or ABC transporters (type II) and export pathways that employ vesicular intermediates (type III) (9). A prominent example for direct translocation is the secretion of mammalian Fibroblast Growth Factor 2 (FGF2), a proangiogenic mitogen, from tumor cells (10). This protein is recruited to the plasma membrane via interaction with phosphatidyl inositol 4,5 phosphate (PI4,5P_2_) (11, 12). At the membrane it assembles into lipidic toroidal pores that surpass the membrane and mediate export of FGF2 monomers (13). Translocation critically depends on phosphorylation via TEC kinase at the membrane. In addition, extracellular heparan sulfates play a crucial role in supporting export by binding FGF2 with high affinity (11, 14). Interestingly, unconventional secretion mechanisms of other important proteins like HIV-Tat or interleukin 1-β share at least some of the features that have been described for FGF2 export (6).

Unconventional secretion has not only been observed in mammalian cells but is also conserved in lower eukaryotes such as fungi. Here, important examples are superoxide dismutase and acyl-CoA binding protein which are secreted by a type III mechanism and have been intensively studied in *Saccharomyces cerevisiae* (15). Both proteins are exported by nutrient-starvation induced unconventional secretion that relies on the Golgi reassembling and stacking protein (GRASP) homolog Grh1 and multiple components of the endosomal sorting complexes required for transport (ESCRT) (16). More specifically, a diacidic motif is exposed upon starvation(17), determining capture of the cargo into Grh1 containing so-called compartments of unconventional secretion (CUPS), that are derived from the Golgi apparatus and from endosomes. The CUPS likely serve as sorting stations for export (15). Unconventional secretion mechanisms of many other proteins are often not fully understood. Nevertheless, this evolving landscape of alternative export pathways highlights the complexity and versatility of cellular mechanisms, expanding our understanding from canonical to unconventional secretion processes.

In *Ustilago maydis*, a cytokinesis-dependent unconventional secretion mechanism was demonstrated crucial for export of the chitinase Cts1 (18, 19). In cylindric yeast cells of this fungal model, cytokinesis is initiated by the formation of a small bud at one of the cell poles which extends by polar growth. After nuclear division, physical separation is initiated by the formation of a primary septum at the mother cell side. The septum acts as a physical barrier and separates the cytoplasm of the two cells. Consecutively, a secondary septum is formed at the daughter cell side, delimiting a membrane-rich so-called fragmentation zone (20). Strains lacking chitinases Cts1 and Cts2 are not able to separate and form tree-like structures, indicating that both enzymes contribute to physical cell separation (21). Interestingly, while Cts2 contains a classical signal peptide this is lacking in Cts1 while it is still active extracellularly (18). We have demonstrated that this enzyme is exported by an unconventional mechanism, that involves Cts1 recruitment from the daughter cell to the primary septum. After formation of the secondary septum, the enzyme is entrapped in the fragmentation zone where it participates in cell separation presumably by hydrolyzing remnant chitin (18, 21, 22).

Recently, we used a forward genetic screen to identify factors involved in unconventional secretion of Cts1. Here, we employed a specifically designed screening strain carrying three reporters for unconventional secretion: intrinsic Cts1, LacZ-Cts1 (fusion to β-galactosidase enzyme) and Gus-Cts1 (fusion to β-glucuronidase enzyme). After UV mutagenesis of the reporter strain, we screened for absence of extracellular Cts1, Gus and LacZ activity and discovered the yet uncharacterized protein Jps1 (22). *jps1* deletion mutants show mis-localized Cts1 that is excluded from the fragmentation zone and a strongly diminished extracellular chitinase activity, supporting its essential role for Cts1 export (22). Jps1 lacks a predicted signal peptide but is present in the culture supernatant, indicating that it is also exported unconventionally (23). In line with that, Jps1 is enriched in the fragmentation zone, as observed for Cts1, and yeast-two hybrid data support the idea of a direct interaction (22). Hence, we hypothesized that Jps1 might be an anchoring factor that recruits or tethers Cts1 to the fragmentation zone. Here, we now present an in-depth biochemical analysis of Jps1, revealing the presence of a core domain that mediates dimerization and an unusual mode of PI(4,5)P_2_ binding. Interestingly, these features are conserved across different orthologs in the basidiomycete phylum. Our findings are in line with the essential function of Jps1 in the membrane-rich fragmentation zone and provide new insights into this mechanism of unconventional secretion.

## Results

### Jps1 is a conserved basidiomycete-specific protein with a central core domain

Our previous work identified Jps1 as an essential factor for unconventional secretion in *U. maydis* (22). No information on the function of Jps1 was available from public databases and domain prediction tools did not reveal any known functional domains. We therefore performed a phylogenetic search using BlastP (https://blast.ncbi.nlm.nih.gov/Blast.cgi) with the 609 amino acids sequence of *U. maydis* Jps1 protein as a query to identify conserved patches that might hint to functionally relevant regions. Orthologs were present and widespread but restricted to basidiomycetes. In total, more than 1,000 homologs of Jps1 were identified across these different basidiomycete species. We observed a large variability in sequence length, ranging from ∼185 to more than 1,000 amino acids. Consistently, a region covering residues 356-480 of the *U. maydis* Jps1 variant, showed an overall high conservation (35-53 % sequence identity) across all species, while the N- and C-terminal regions had a high degree of variability in the amino acid sequence and length (**Fig. S1**). Notably, the conserved region is also present in short Jps1 versions, as e.g. the one of *Hebeloma cylindrosporum* that only consists of 287 amino acids (**Fig. S1**).

To gain a better understanding of the architecture of Jps1, we used AlphaFold2 and predicted a structural model of the protein (**Fig. S2**). As we did not have prior information of the oligomeric state of Jps1, we also predicted homodimers, trimers and tetramers using AlphaFold2-Multimer (24). The Jps1 monomer showed two regions with intermediate pLDDT values (>80), while the aforementioned conserved region had high pLDDT values of >90 (**Fig. S2A**). Predicting a Jps1 dimer elevated the pLDDT in the conserved regions from around 70-80 to >95, while it strongly dropped in the less conserved regions supporting the presence of intrinsically disordered regions (**Fig. S2A, B**). Prediction of higher oligomers did not improve the pLDDT as observed for the dimer (data not shown). Our structural model suggests that Jps1 has unstructured N- and C-termini and a central domain that is assembled from several structured parts within the N-terminus and the conserved core domain (**Fig. 1A, B**). This core domain consists of 8 β-strands and a total of 11 α-helices (**Fig. 1B**). Six of the β-strands (β1, 2, 5-8) form a central β-sheet that is framed by helices α1-4, 7 and helices α10-12 on the other side (**Fig. 1B**). Helices α8, 9 and the two β-strands 3 and 4 form a small domain that is stacked onto the central β-sheet (**Fig. 1B**). This central domain is apparently not only formed by the above-mentioned core region of Jps1 but residues located N-terminally also contribute (e.g. α1-α7 and β1, β2 and β3). Three flexible loop regions (LR) surround the compact protein core (LR1-LR3, **Fig. 1A, B**).

**Figure 1:**
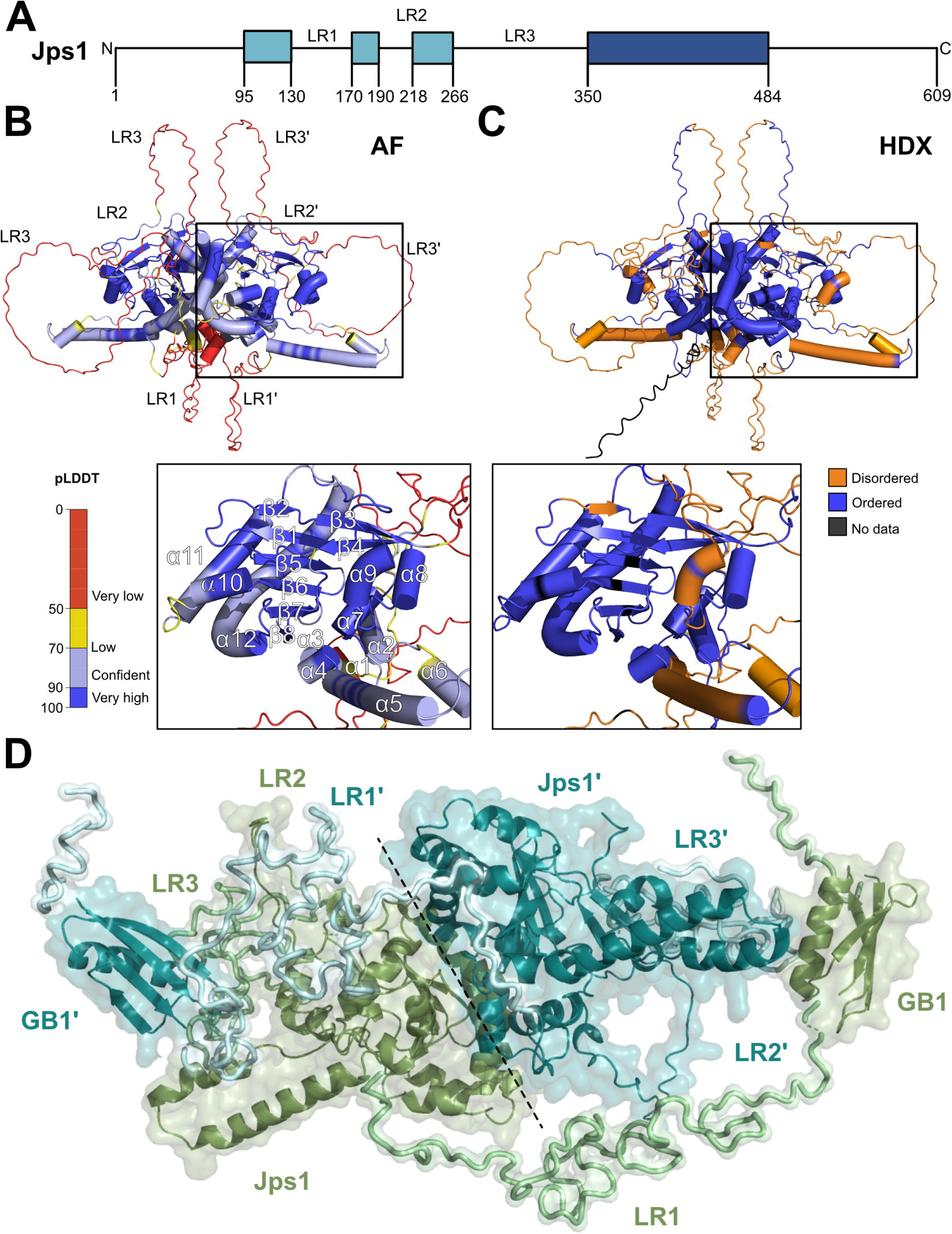
Structure prediction and HDX-MS reveal a flexible architecture of Jps1. **A.** Domain architecture of Jps1 based on the structural prediction (AF) and HDX-MS analysis. Boxes indicate structured patches, while the coloring reflects less conserved regions (light blue) and the region strongly conserved across basidiomycetes (dark blue). LR: Loop regions. **B.** Structural model of the Jps1 dimer predicted with AlphaFold2 (AF) (24) shown in cartoon representation and colored according to the model confidence. **C.** Structural model of the Jps1 dimer colored according to HDX-MS exchange. Disordered and ordered regions and those not covered in HDX-MS (HDX) are colored in red, blue and black in the AlphaFold2 model of Jps1, respectively. Below the two panels, a close-up of the central core domain is shown. Secondary structure elements are labeled accordingly. **D.** Rigid-body modeling of the Jps1 dimer based on small-angle X-ray scattering data. The two protomers, Jps1 and Jps1’, are colored in smudge green and cyan, respectively.

To support our bioinformatic analysis on Jps1 and gain information on its biochemical properties, we heterologously produced Jps1 as GB1 fusion in *Escherichia coli* and purified the soluble protein through a two-step purification procedure consisting of Ni-NTA affinity chromatography followed by size-exclusion chromatography (**Fig. S3A**). We subjected the purified Jps1 to hydrogen-deuterium exchange mass spectrometry (HDX-MS) to validate the structured and flexible regions of Jps1 suggested by our structural model. Mapping our HDX-MS results revealed that most of the Jps1 residues inferred as disordered in the AlphaFold2 model reach their maximal deuterium incorporation already after ten seconds of HDX while those constituting the core domain exhibit protection against fast HDX indicative for the presence of secondary or higher-order structures (**Figs. 1C, S4**). This suggests that Jps1 adopts a conformation that is reminiscent of its structural model in solution. To further confirm these findings, we also performed small-angle X-ray scattering (SAXS) experiments. Here, we generated a Jps1 version lacking the unstructured C-terminal residues (Jps1^1-484^) as this region does not contribute to the structured core domain and a large number of disordered residues complicates the *ab initio* modeling. Our SAXS experiments confirmed an overall compact architecture of the Jps1 dimer with some elongated regions as judged from the intra-particle distance and the Kratky plot (**Fig S5, Table S1**). We also performed rigid body modeling of Jps1, which suggested that the loop regions (**Fig. 1B**) most likely align more closely to the protein core (**Fig. 1D**). In conclusion, a combination of bioinformatic and biochemical experiments revealed that Jps1 is a basidiomycete-specific protein with a core domain enclosed by flexible regions.

### Jps1 dimerizes via the core domain

To further characterize Jps1 and confirm the predicted dimer, we performed size-exclusion chromatography coupled multi-angle light scattering (SEC-MALS). Jps1 eluted in one stable fraction from the SEC, while MALS revealed the presence of a single species of 134 ± 3.5 kDa indicating the formation of homodimers based on the calculated molecular weight (MW) of 63 kDa of a Jps1 monomer (**Fig. 2A**). In addition, we also performed mass photometry experiments using a nanomolar concentration (10 nM) of Jps1 showing that 69% of all particles were in a single fraction at 125 kDa matching the MW of dimers, with only a small subfraction of 9% at 64 kDa (**Fig. 2B**). This is in line with the SAXS experiments that also revealed a predominantly dimeric species in solution (**Tab. S1**).

**Figure 2:**
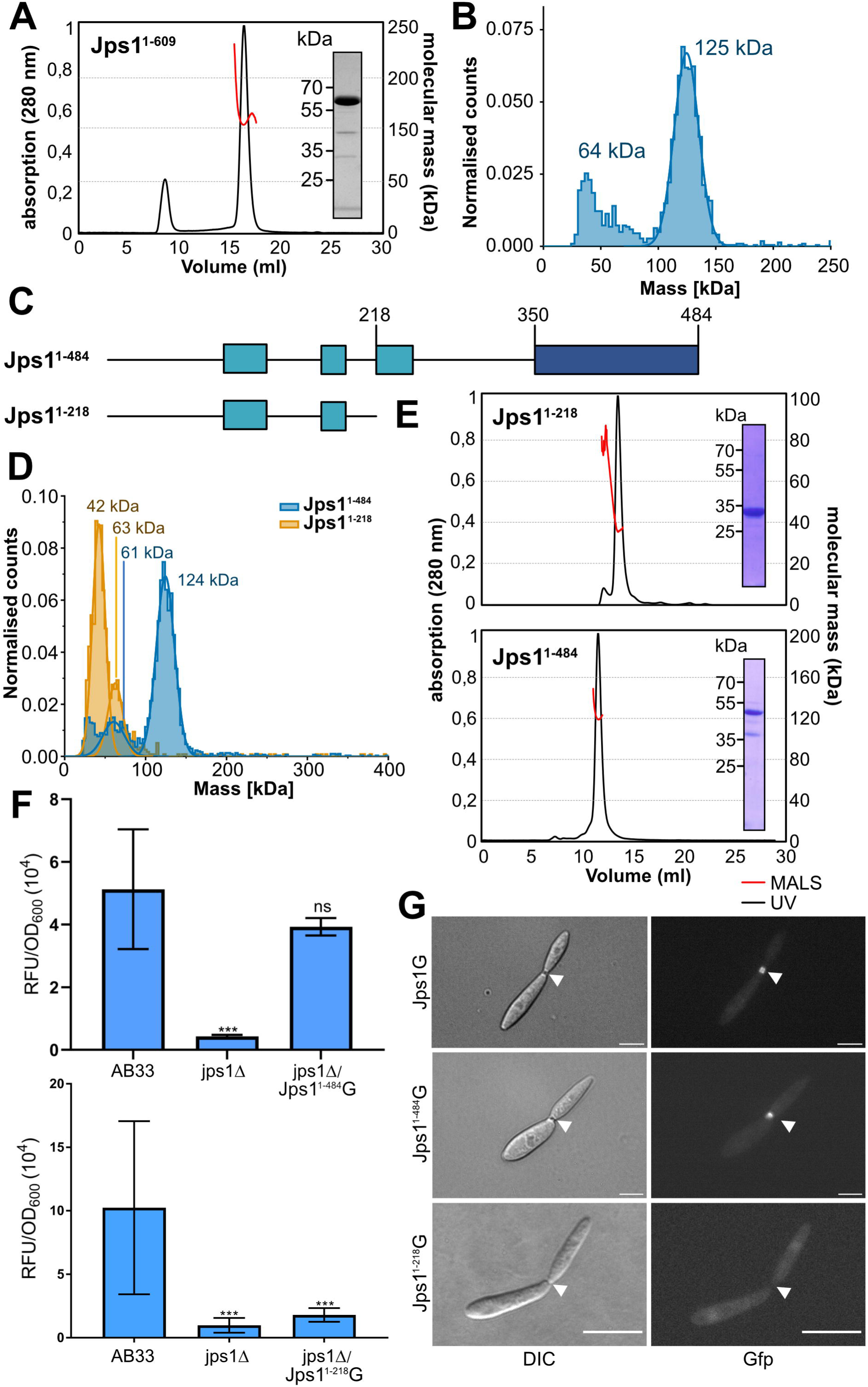
Biochemical analysis of *U. maydis* Jps1 supports homodimer formation. **A.** Multi-angle-light scattering coupled SEC (SEC-MALS) of full length recombinant UmJps1 (Jps1^1-609^). The black line shows the absorption at 280 nm (SEC), the molecular weight as determined by MALS is depicted in red. Inset: SDS-PAGE of purified protein before SEC-MALS. **B.** Mass photometry of recombinant UmJps1 at a concentration of 10 nM. **C.** Schematic representation of the protein architecture of the two truncated UmJps1 versions Jps1^1-484^ and Jps1^1-218^. Boxes indicate structured patches, while the coloring reflects less conserved regions (light blue) and the region strongly conserved across basidiomycetes (dark blue). **D.** Mass photometry of the two variants UmJps1^1-484^ and UmJps1^1-218^ at a concentration of 10 nM. **E.** SEC-MALS of UmJps1^1-484^ and UmJps1^1-218^. The black line shows the absorption at 280 nm (SEC), the molecular weight as determined by MALS is depicted in red. Insets: SDS-PAGE of purified proteins before SEC-MALS. **F.** Extracellular Cts1 activity of indicated AB33 derivatives. Error bars depict standard deviation. ***P-value 0.001; n.s., not significant (two sample t-test). The assay was conducted in three biological replicates. **G.** Fluorescence microscopic localization of Jps1G and the truncated version Jps1^1-484^G in yeast-like growing cells AB33 jps1Δ derivatives (25 and 25 cells examined, respectively). DIC, Differential interference contrast. Scale bars, 10 µm.

To identify the region of Jps1 that mediates the homodimerization, we tested the previously mentioned truncated version lacking the unstructured C-terminus (Jps1^1-484^, 60 kDa) and an N-terminal version lacking the conserved domain (Jps1^1-218^, 32 kDa)(**Fig. 2C**), again fused to an N-terminal GB1-tag. Both truncated Jps1 constructs could be expressed and purified to homogeneity (**Fig. S6A**) and employed for mass photometry measurements. Experiments on Jps1^1-218^ revealed a dominant fraction at 42 kDa and a small subfraction at 63 kDa, while for Jps1^1-484^ similar results as for the full-length Jps1 were observed (**Fig. 2D**). MALS of the purified variants also indicated a molecular weight of 36.5 ± 1.9 kDa of Jps1^1-218^ suggesting a monomer while Jps1^1-484^ formed dimers comparable to the full-length protein with a molecular mass of 117.2 ± 1.1 kDa (**Fig. 2E**).

Earlier results from the forward genetic screen demonstrate that three Jps1 versions that are truncated by insertion of a premature stop codon within the conserved core domain are not functional (22). To assess whether a protein version with the complete core domain but lacking the adjacent flexible C-terminal region is still functional, we expressed the construct for Jps1^1-484^ fused to Gfp (Jps1^1-484^G) in a *jps1* deletion strain (AB33 jps1Δ/Jps1^1-484^G). *In vivo* studies revealed full complementation of extracellular Cts1 activity, suggesting that the truncated protein is fully functional in mediating unconventional Cts1 secretion (**Fig. 2F**). In accordance, fluorescence microscopic inspection of the respective strains confirmed the localization of Jps1^1-484^G in the fragmentation zone of dividing cells (**Fig. 2G**). By contrast, a complementation strain expressing the Jps1 version truncated for the core domain (Jps1^1-218^G) did show ablated extracellular Cts1 activity and does not localize to the fragmentation zone (**Fig. 2G**). In essence, the conserved C-terminal domain of Jps1 mediates homodimerization and dimerization is crucial for protein function while the adjacent flexible C-terminal region is dispensable.

### Jps1 orthologs homodimerize but exhibit functional diversification

Our phylogenetic analysis suggested that the core domain is conserved across fungal species, while the surrounding regions show a high sequence variability (**Fig. S1**). We were therefore interested whether dimerization is a conserved feature of Jps1 and whether orthologs can functionally complement the *jps1* deletion. Two Jps1 orthologs were selected for recombinant production in *E. coli*: A long version from the very close relative *Sporisorium reilianum* (Sorghum smut, SrJps1; calculated MW of 77 kDa) and a short Jps1 version from the mushroom *Hebeloma cylindrosporum* (HcJps1; 38 kDa). To avoid effects of codon bias in complementation studies, we dicodon-optimized the gene sequence for HcJps1 to match the codon preferences of *U. maydis*. Both versions were well-expressed as soluble GB1-fusions and could be purified as described previously (**Fig. S6B**). SEC-MALS revealed that SrJps1 and HcJps1 constituted dimers with apparent molecular masses of 153.2 ± 2.7 kDa and 77.8 ± 0.6 kDa, respectively (**Fig. 3A**). Notably, mass photometry suggested that SrJps1 formed predominantly dimers, while we observed mostly monomeric species of HcJps1 (**Fig. 3B**). Our experiments therefore suggest that the HcJps1 ortholog is able to adopt both a mono- and a dimeric state, which likely depend on the protein concentration. To assess whether the conserved core domain also forms a similar structural fold in SrJps1 and HcJps1, we predicted the structural models using AlphaFold2. Overall, the models share the architecture of UmJps1 and superpose well with RMSD of 0.8 (SrJps1) and 1.4 (HcJps1) over 287 and 136 Cα-atoms, respectively, within the central core domain (**Fig. S7**).

**Figure 3:**
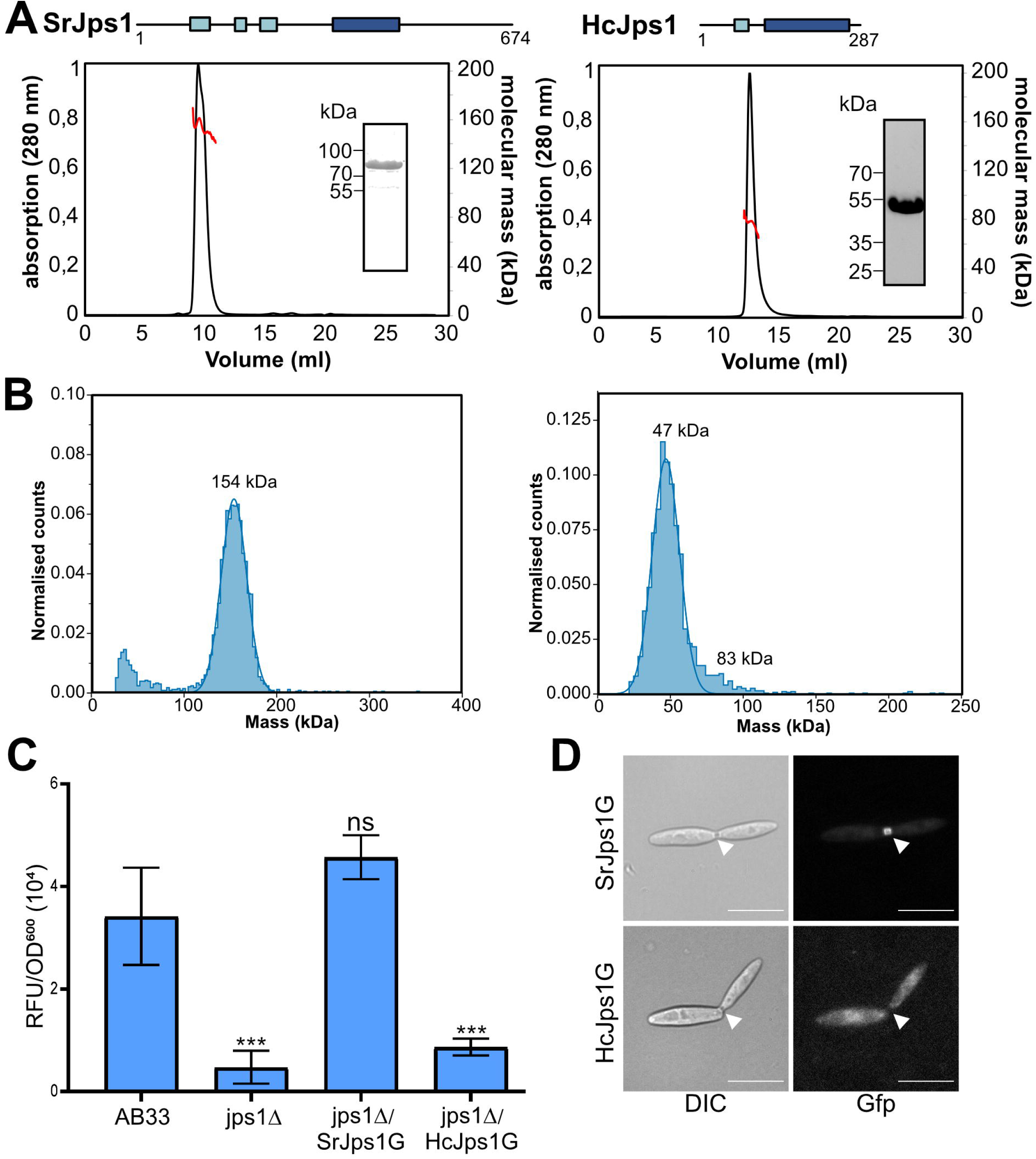
Biochemical analysis of Jps1 orthologs confirms dimerization but reveals functional diversification. **A.** Schematic representation of SrJps1 (left) and HcJps1 (right) protein architecture. The conserved core domain is depicted in dark blue. Below, SEC-MALS results of full-length recombinant SrJps1 and HcJps1 are shown. The black line indicates the absorption at 280 nm (SEC), the molecular weight as determined by MALS is depicted in red. The insets show SDS-PAGE of purified proteins before SEC-MALS. **B.** Mass photometry of SrJps1 (left) and HcJps1 (right) at 10 nM concentration. **C.** Extracellular Cts1 activity of indicated strains. AB33jps1Δ was used as negative control, the progenitor AB33 dealt as positive control. Error bars depict standard deviation. ***P-value 0.001; n.s., not significant (two sample t-test). The assay was conducted in three biological replicates. **D.** Fluorescence microscopic localization of SrJps1-G and HcJps1-G in the complementation strains (24 and 20 cells examined, respectively). DIC, differential interference contrast. Scale bars, 10 µm.

To analyze the functional conservation of the orthologs *in vivo*, we generated complementation strains in which we produced the protein versions as translational fusions with Gfp (SrJps1G, HcJps1G) in the *jps1* deletion strain. Both fusion proteins were produced in full-length as verified by western blotting (**Fig. S8A**). Biochemical *in vivo* studies revealed full complementation of extracellular Cts1 activity for the strain expressing SrJps1G (**Fig. 3C**). By contrast, no complementation was achieved by the short version HcJps1, and extracellular chitinase activity could not be detected (**Fig. 3C, S8**). Instead, intracellular Cts1 activity was increased in strains producing HcJps1G, suggesting that the protein remained and accumulated in the cells as observed for *jps1* deletion strains (**Fig. S8B**). Accordingly, fluorescence microscopy using the respective strains revealed a localization of SrJps1G but not HcJps1 in the fragmentation zone (**Fig. 3D**). In summary, the characterization of Jps1 orthologs confirmed the conservation of the dimerization via the core domain. Our complementation studies further suggest that Jps1 has functionally diversified in more distantly related basidiomycete species, resulting in the correct subcellular localization of the *S. reilianum* ortholog but not the one from *H. cylindrosporum*.

### Jps1 interacts with phosphatidylinositol phosphates

Jps1 localizes in the membrane-rich fragmentation zone and particularly accumulates at the membrane enclosing the small compartment which is likely derived from the plasma membrane (19, 25). This membrane localization could either be mediated through a direct lipid interaction or indirect via interaction with a yet unknown factor. To test these two scenarios, we again used purified recombinant Jps1 to screen for membrane affinity using commercially available PIP strips (Echelon Biosciences Inc.) in lipid-protein interaction assays. Interactions of varying strengths were detected when Jps1 was incubated with the PIP strip membrane, including all phosphatidylinositol phosphates (PIP) and phosphatidic acid (PA; **Fig. S9A**). To corroborate this membrane binding, we generated heavy liposomes containing DOPC, the varying phospholipids or PA, and eventually cholesterol and performed interaction assays by mixing the liposomes with recombinant protein followed by high-speed centrifugation, initially using the full range of PIPs and PA shown positive in the PIP strip assay. The recombinant NADPH oxidase p40 (PX) domain of *U. maydis* Yup1 (26) was used as a positive control known to bind PI3P (phox-Yup1; **Fig. S9B**). While Jps1 did not bind to pure DOPC liposomes or liposomes containing PA, we detected weak binding to all PIPs except for PI(3,4,5)P_3_ and to cholesterol. In some cases, like for PI(3,4)P_2_ and PI(5)P the addition of cholesterol slightly increased the binding to the PIPs (**Fig. S9C**). Overall, we did not observe strong PIP binding and also noted degradation of Jps1 during our assays. We therefore decided to test, whether the previously characterized orthologs SrJps1 and HcJps1 could be used for the liposome assays instead. SrJps1 behaved similarly to UmJps1 but showed stronger binding to PI(4,5)P_2_ compared to the other used PIPs (**Fig. S9D**). A substantial subfraction of HcJps1 bound to PI(4,5)P_2_ liposomes in both the absence and even better in the presence of cholesterol. In the case of PI(4)P, a slightly weaker binding to the liposomes was detected (**Fig. S9E**). Overall, our results therefore suggested that HcJps1 preferentially binds to PI(4,5)P_2_, while instability of UmJps1 and SrJps1 resulted in a less stringent binding behavior. To characterize PI(4,5)P_2_ interaction in more detail, we repeated the liposome binding assays for this PIP with all three proteins in biological triplicates. In addition, we included truncated protein versions of Jps1, either including (Jps1^1-484^) or excluding the core domain (Jps1^1-218^) (**Fig. S10**). Strong binding was confirmed for HcJps1 with about 50% of the input protein sticking to the liposomes, rising to about 75% binding in the presence of cholesterol. Moderate binding was observed for UmJps1 and SrJps1 with approximately 50% binding in the presence of cholesterol. Interestingly, in the truncated variant UmJps1^1-484^ liposome binding increased to a level comparable to HcJps1. By contrast, elimination of the core domain in UmJps1^1-218^ also ablated PI(4,5)P_2_ interaction (**Fig. S10**). For further verification, we generated giant unilamellar vesicles (GUVs) containing cholesterol and either PI(4,5)P_2_ or PI(3,4,5)P_3_. To enable microscopic inspection, we generated recombinant UmJps1 fused to Gfp (Jps1G) and Gfp as a negative control in *E. coli*. Interestingly, UmJps1G showed enhanced PIP binding with a clear preference to PI(4,5)P_2_ as compared to UmJps1 in liposome assays (**Fig. S9F**). Next, the purified proteins were incubated with the different GUVs and interaction was visualized by fluorescence microscopy. In these assays, no interaction was detected for PI(3,4,5)P_3_, while a strong and stable association of Jps1G was obtained for PI(4,5)P_2_ containing GUVs (**Fig. 4A**). Hence, strong experimental evidence suggests that PI(4,5)P_2_ is the major PIP interacting with Jps1.

**Figure 4:**
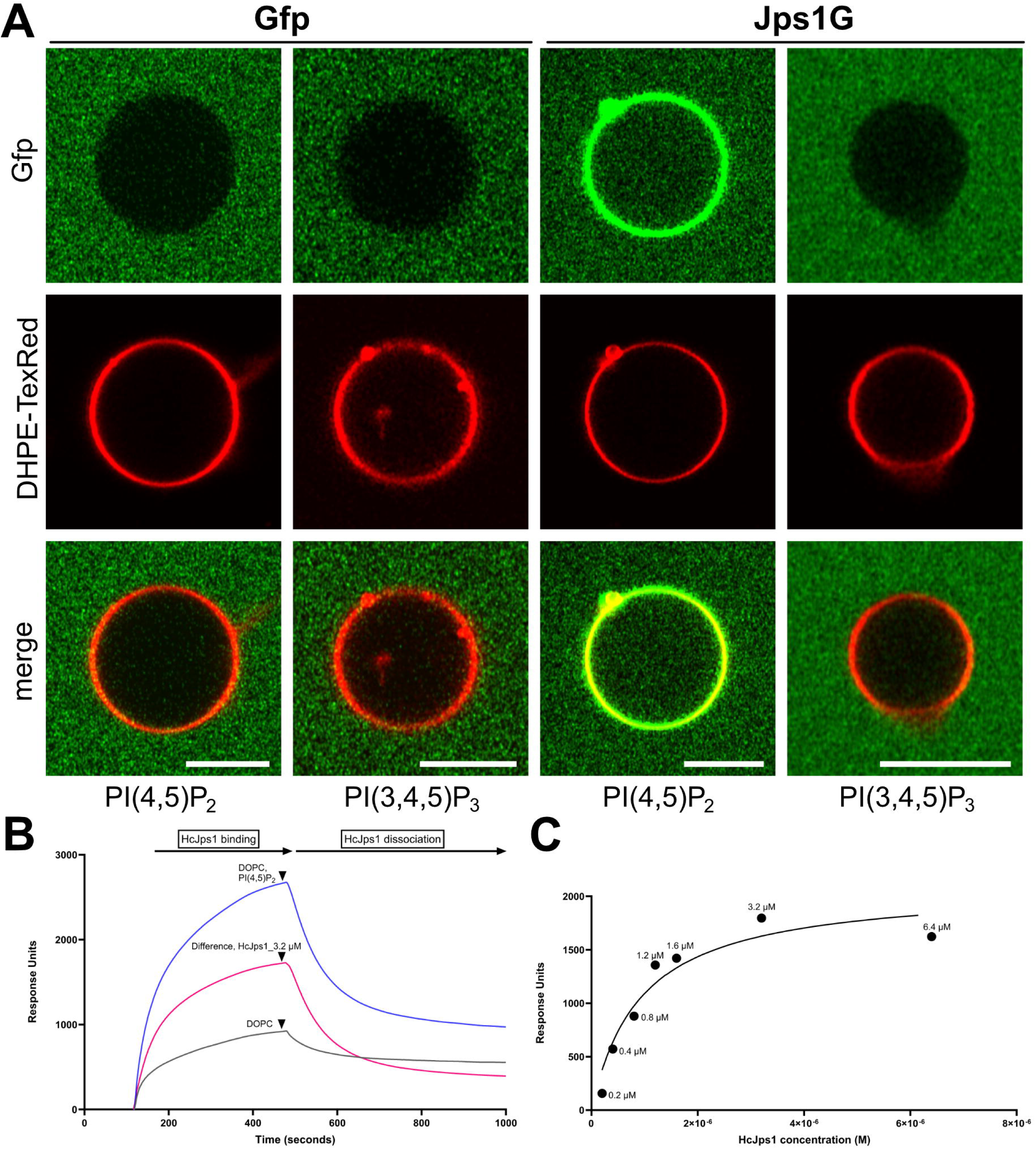
GUV- and liposome-binding studies verify Jps1 affinity to PI(4,5)P_2_. **A.** Microscopic visualization of interaction of Jps1 with GUVs. GUVs containing the indicated PIPs were incubated with recombinant Jps1 fused to Gfp (Jps1G). Recombinant Gfp was used as a negative control, DHPE-TexRed was employed to stain vesicle membranes. **B.** Surface Plasmon Resonance (SPR) of HcJps1 at 3.2 µM indicating binding to the test liposomes DOPC, PI(4,5)P_2_ shown as the difference between the test and the reference channel (DOPC-only liposomes). **C**. Analysis of the SPR response units of HcJps1 at concentration range of 0.2 to 6.4 µM binding to immobilized liposomes fitted by non-linear regression using the steady-state affinity model. Liposomes containing DOPC and PI(4,5)P_2_ at 1 mM concentration were used. DOPC-only liposomes were used a negative control in the reference channel.

To further characterize phosphoinositide binding, we performed quantitative analyses by surface plasmon resonance spectroscopy (SPR) using hydrophobic sensor chips with attached PI(4,5)P_2_^-^containing liposomes and different concentrations of purified Jps1. While the interaction of UmJps1 with liposomes was difficult to address likely due to aggregation on the chip, we could confirm the specific interaction of HcJps1 with PI(4,5)P_2_ containing liposomes (1 mM) at concentrations between 0.2 and 6.4 µM, after which the response units reached the saturation level (**Figs. 4B, C, S11**). The measurements obtained in this concentration range provided an estimate of the dissociation constant K_D_ for HcJps1:PI(4,5)P_2_ interactions to be 0.9 µM (900 nM). This notably signifies that HcJps1 displays a strong binding affinity to PI(4,5)P_2_. In essence, Jps1 displays PIP binding activity that likely mediates membrane attachment with specificity for PI(4,5)P_2_-rich membranes.

### Disturbing PIP-binding specificity causes morphological perturbations

To delineate the details of Jps1:PI(4,5)P_2_ interaction and its implications for unconventional secretion we next focused on identifying regions in the protein that mediate PIP interaction. Due to the absence of a canonical PIP binding domain, we hypothesized that basic residues are involved and selected lysine/arginine (K/R) clusters in the N-terminal part of Jps1^1-484^ to generate 5 mutagenized versions with alanine (A) replacements for recombinant expression in *E. coli* (**Fig. S12A**). Protein versions Jps1^1-484^M^1^, -M^3^, -M^5^ were soluble and could be purified to homogeneity (**Fig. S12B**) while Jps1^1-484^M^2^ and M^4^ were not expressed in sufficient amounts, suggesting that these replacements might have destroyed essential structural interactions. Unexpectedly, liposome binding studies did not reveal any reduction of PIP binding in the three variants (**Fig. S12C**). However, we observed that Jps1^1-484^M^5^ has lost its PIP binding specificity and now showed interaction also with liposomes containing only DOPC or DOPC and cholesterol (**Fig. 5A**). Notably, MALS analysis of Jps1^1-484^M^5^ revealed the retained presence of dimer conformation with a MW of 118.3 ± 0.025 kDa (**Fig S12D**).

**Figure 5:**
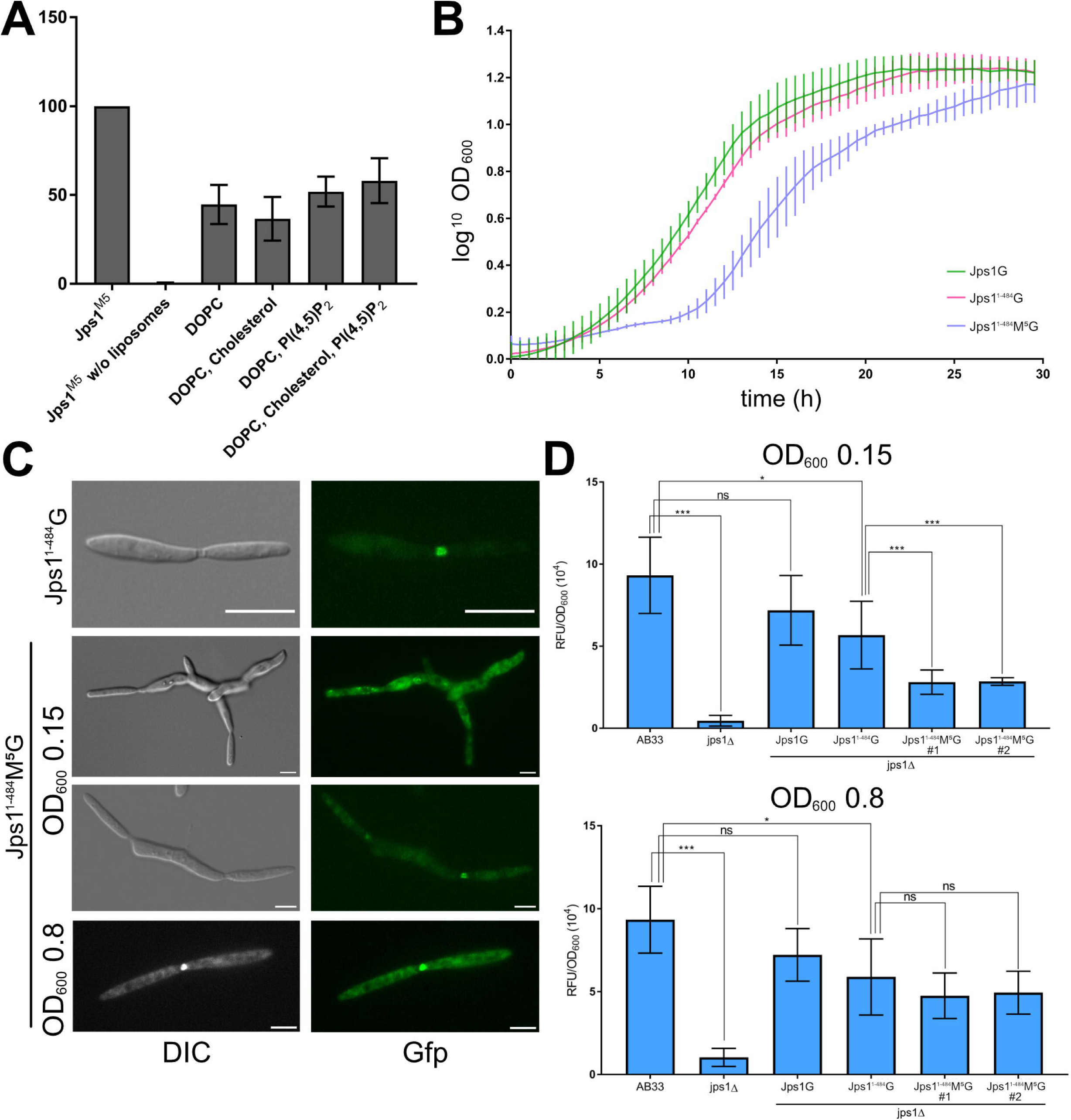
Mutation of a basic cluster results in loss of PIP specificity and causes defects in unconventional secretion, growth and morphology. **A**. Quantitative evaluation of liposome binding assays using liposomes containing PI(4,5)P_2_ and the protein variant Jps1^1-484^M^5^. The assays were conducted in three replicates. Error bars depict standard deviation. **B.** Growth curves of the indicated strains obtained from BioLector micro-cultivations. Growth was followed online using scattered light measurements. The growth assays were conducted in three biological triplicates. Error bars represent standard deviation. **C**. Fluorescence microscopy visualizing cell morphology and the localization of Jps1^1-484^G and Jps1^1-484^M^5^G in complementation strains. Scale bars, 10 µm. **D**. Extracellular chitinase activity assayed for the indicated strains at OD_600_ of 0.15 (lag phase) and 0.8 (exponential growth phase). The assays were conducted in three biological replicates. Error bars depict standard deviation. ***P-value 0.001; *P-value 0.05; n.s., not significant (two sample t-test).

To check the functional consequences of the M^5^ mutation *in vivo*, we next used a Gfp fusion variant (Jps1^1-484^M^5^G) for complementation of the deletion mutant (AB33 jps1Δ/jps1^1-484^M^5^G). Growth curves recorded in a micro-cultivation device revealed that the strain producing the mutant version exhibited an extended lag phase of about 5 h as compared to the control strains complemented with full length Jps1G and the truncated version Jps1^1-484^G (AB33 jps1Δ/jps1G and AB33 jps1Δ/jps1^1-484^G) while the growth rate during exponential phase was unchanged at a generation time of approximately 2.3 h (**Fig. 5B**). In line with this, microscopic inspections unveiled morphological perturbations with elongated cells showing irregular shapes and partially cytokinesis defects. Intriguingly, these morphological defects were specifically detected at low cell densities (OD_600_ 0.15) while cells gradually returned to a normal shape at higher cell densities (OD_600_ 0.8) (**Fig. 5C**). Fluorescence imaging uncovered that at low optical density Jps1^1-484^M^5^G in comparison to the control was enriched in the cytoplasm and in larger cytoplasmic accumulations (**Fig. 5C**). Septal staining revealed that fragmentation zones with a Jps1^1-484^M^5^G signal were mostly formed in these cells although cytokinesis and cell morphology were disturbed (**Fig. S13**). Chitinase assays confirmed that extracellular Cts1 activity was strongly reduced at low cell densities in the strain producing the M^5^ mutant variant as compared to the strain containing Jps1^1-484^G while it mostly recovered at high densities (**Fig. 5D**). In summary, PI(4,5)P_2_ binding specificity is crucial for Jps1 localization and unconventional Cts1 secretion at low optical densities and its disturbance results in unexpected cellular defects. Thus, we provide strong evidence that Jps1 dimers are targeted to the zone of unconventional secretion by a direct interaction with PI(4,5)P_2_, thereby facilitating unconventional secretion of Cts1.

## Discussion

In recent years, different unconventional secretion mechanisms have been described in eukaryotes that deliver a variety of target cargo to the plasma membrane and beyond (6–8). We have previously reported a new type of unconventional secretion mechanism underlying export of the chitinase Cts1 in *Ustilago maydis* (18, 19). We furthermore identified the protein Jps1 as important factor for Cts1 secretion while lacking detailed insights (22, 23). Here, we therefore combined *in vitro* experiments with recombinant protein and *in vivo* studies to investigate how Jps1 contributes to this export mechanism. Our findings reveal that Jps1 directly targets the membrane through an intrinsic affinity to PIPs, particularly to PI(4,5)P_2_. A combination of biochemical and biophysical techniques furthermore demonstrated that Jps1 is a flexible and highly dynamic protein that homodimerizes through a conserved structural domain. Thus, our findings not only allow to refine the model of unconventional secretion of Cts1 in *U. maydis* but also expand our current knowledge on unconventional secretion mechanisms.

### Expanding the model of unconventional secretion in *Ustilago maydis* and beyond

The recruitment of homodimeric Jps1 to the fragmentation zone, likely mediated by interaction with PI(4,5)P_2_, a PIP enriched in the inner leaflet of the plasma membrane, supports the hypothesis that Jps1 serves as a tether for Cts1 accumulation at the site of cell separation (**Fig. 6**). In line with that, we see a slight interaction of Jps1 and Cts1 in yeast-two hybrid assays and Cts1 is excluded from the fragmentation zone in the absence of Jps1 (22). The precise Cts1 interaction interface at Jps1 has not been clarified yet. However, the expanded structural architecture of the Jps1 homodimer offers ample possibilities for protein:protein interactions. Truncating the conserved core domain of Jps1 diminishes extracellular chitinase activity (22), suggesting that Cts1 interaction is either mediated by this part of the protein or homodimerization is a prerequisite for proper function of Jps1. The process of unconventional Cts1 secretion might also involve other factors that interact with Jps1 and require the presence of the structural domains. Such protein:protein interactions could also be critical to determine the specificity of binding PIPs in the fragmentation zone but not in the mother or daughter cell. This hypothesis is further supported by the poor sequence conservation in Jps1 orthologs apart from the core domain.

**Figure 6:**
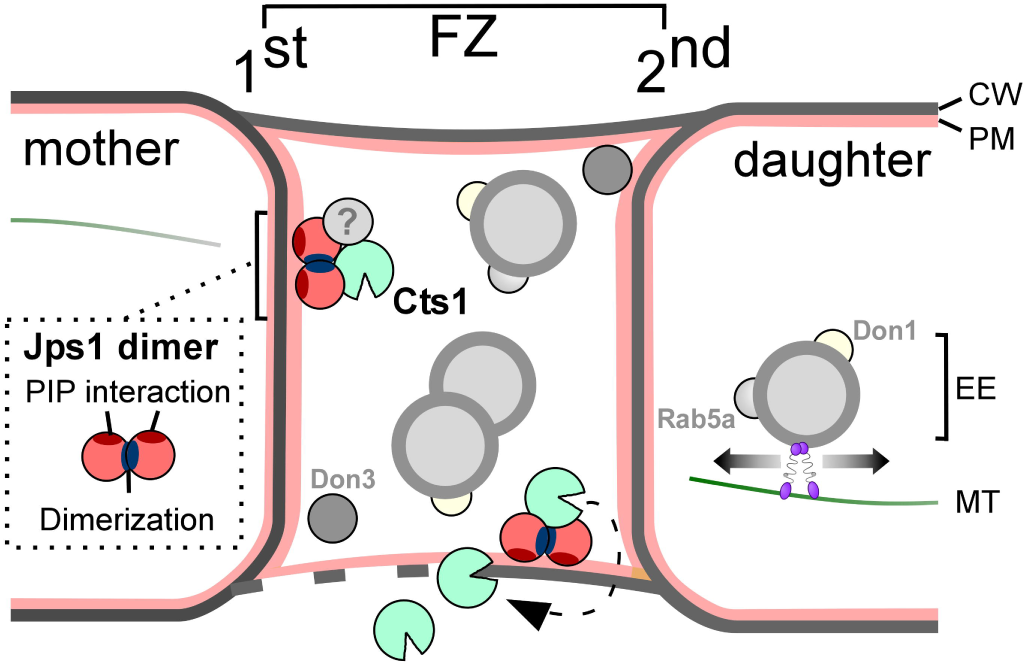
Schematic model of cytokinesis-dependent unconventional secretion. The picture shows the fragmentation zone (FZ) between two dividing yeast cells that is delimited by the primary (1^st^) and secondary septum (2^nd^). The fragmentation zone is completely encased by a peripheral membrane that is likely derived from the plasma membrane (PM). Jps1 and Cts1 both accumulate in the fragmentation zone. The former acts as a (homo)dimer and presumably interacts with the PIP PI(4,5)P_2_ of the peripheral membrane. We speculate that specificity for the FZ membrane is mediated by yet unknown interaction partners (indicated by ?). Our data support the hypothesis that Jps1 recruits chitinase Cts1 to the FZ. After secretion, the chitinase supports cell division by hydrolyzing remnant chitin. The fragmentation zone also contains early endosomes (EE) that shuttle bidirectionally throughout the yeast cells and accumulate in the FZ. Motile EE carry the proteins Rab5a and the GEF Don1. Kinase Don3 resides in the FZ. CW, cell wall; MT, microtubules.

In the course of this investigation, we therefore also explored a potential functional conservation of Jps1 orthologs within the phylum of basidiomycetes. Jps1 of *S. reilianum*, a very close dimorphic relative and a pathogen of corn and sorghum (27), shows high sequence conservation as well as complete functional complementation and a similar biochemical behavior. By contrast, the other – more distant – ortholog from *H. cylindrosporum* shares biochemical key features like dimerization and PI(4,5)P_2_ affinity while functional complementation *in vivo* is lost. The strong reduction in protein size for this ortholog suggests that the protein has undergone evolutionary adaptations to its lifestyle. Interestingly, *H. cylindrosporum* is a mushroom-forming, hyphal fungus and yeast-like growing stages are not described (28). In the filamentous form of *U. maydis*, Cts1 localizes to the growth apex as well as to retraction septa and the cell wall of empty sections, suggesting an involvement in degradation of remnant chitin in dead hyphal parts (29). Septal localization is in line with the idea of septum-directed secretion described for *Aspergillus oryzae* (30), where classical secretion via exocytosis occurs not only towards the cell exterior but also towards the septal membrane. We speculate that this phenomenon is not restricted to proteins with a canonical signal peptide but also occurs in alternative secretion routes. In this respect it is interesting to note that a glycoside hydrolase 18 (GH18) domain chitinase lacking a predicted signal peptide is present also in *H. cylindrosporum*. However, future research will have to address whether a similar mechanism of unconventional secretion exists in *H. cylindrosporum*. In conclusion, we demonstrate that the specific interaction of Jps1 with PI(4,5)P_2_ is crucial for unconventional secretion of Cts1 in *U. maydis*. PIP binding is conserved across basidiomycete Jps1 orthologs, although their precise functional roles might have diversified.

### Direct interaction with membranes via PIP binding is crucial for unconventional secretion

Multiple experimental evidence has demonstrated that Jps1 has an affinity to PIPs, particularly to PI(4,5)P_2_. This phosphoinositide mainly localizes to the cytoplasmic leaflet of the plasma membrane dealing as a landmark for recruitment of cytosolic proteins and involved in regulating the activity of integral membrane proteins (31). It is crucial for important cellular processes including membrane fusion, signal transduction and also protein translocation (32). Various phosphoinositide binding protein domains have been described in literature, including Pleckstrin-homology domain (PH) mediating binding to various PIPs, Phox-homology (PX) and FYVE zinc finger domains interacting specifically with PI3P (33). All of these domains have a conserved three-dimensional fold that provides the structural scaffold for residues recognizing the phosphate moieties of different phosphoinositides. In Jps1 we did not detect obvious similarities to any of the described phosphoinositide binding domains, suggesting the absence of a structured binding domain. However, besides distinct domains also polybasic stretches can support PIP binding via electrostatic interactions (34). Exploiting mutagenized Jps1 versions we demonstrate that a cluster of basic, positively charged residues (R162, R163, R164) in a variable part of the N-terminus of Jps1 determines PI(4,5)P_2_ binding specificity. Although two versions could not be expressed in *E. coli* and we did not detect changes in PIP binding for the other variants, it might still be possible that a combination of scattered residues could have a detrimental effect on PIP binding. In such case, interactions in the folded protein might bring these clusters in proximity. Although Jps1 has several regions of high flexibility, our SAXS analysis suggests that they rather adopt a compact conformation and align to the central core domain (**Fig. 1D**). We therefore consider it possible that PIP-binding is mediated by residues within these flexible regions, which might even become more rigid and adopt a stable fold in the presence of a bound PIP.

The interaction with PIPs, particularly with PI(4,5)P_2_, is a phenomenon also observed in distinct other pathways of unconventional secretion where proteins are secreted via direct translocation (type I unconventional protein secretion). The detailed insights available for the export of the mammalian FGF2, for example, demonstrate that the protein oligomerizes at the inner leaflet of the cytoplasmic membrane via PI(4,5)P_2_ interaction and this process is essential for secretion (11, 35). Interestingly, also in this case a canonical PIP binding domain is absent while a stretch of three basic residues exposed at the protein surface is responsible for PI(4,5)P_2_ interaction (35). Isothermal titration calorimetry measurements showed a binding affinity with a *K*_D_ of about 0.5 µM for the PI(1,4,5)_3_ head group, which is in the molar range of our results obtained in SPR measurements with recombinant HcJps1. Similar export mechanisms were proposed for other important mammalian proteins like IL-1β or Tau and viral HIV-Tat, and all depend on membrane recruitment via PI(4,5)P_2_ (13, 36, 37). Further proteins that are unconventionally secreted and rely on PI(4,5)P_2_ binding for export are homeodomain proteins like Engrailed-2 (EN2), acting as important spatial determinants of body plan development in humans. Similar to our observations with Jps1, a supporting role of cholesterol for interaction with PI(4,5)P_2_ has also been described for FGF2 and EN-2, however, the underlying mechanistic details are not yet understood (12). Thus, albeit the molecular mechanism of membrane translocation is completely unknown for Jps1, its intriguing to note that PI(4,5)P_2_ and cholesterol binding seem to constitute a key feature that is conserved between unconventional secretion pathways of yeast and mammalian cells.

We did not detect any obvious morphological phenotype in *jps1* deletion mutants (22). Now, disturbing the PIP binding specificity of Jps1 led to an unexpected cellular defect with impaired cytokinesis, abnormal cell morphologies and reduced extracellular Cts1 activity at low cell densities. Typically, the inoculation of a culture in fresh medium results in a lag phase in which the cells are thought to prepare for the coming exponential growth phase by metabolic adaptations (38). Eventually, but not always, cellular stress can result in an extended lag phase (39). It is conceivable that the mis-placement of Jps1 or one of its dragged-along interaction partners at a wrong subcellular site during lag phase is causative for the morphological defects. This new finding now places its cellular role into a new light. Based on the observed lag phase phenotype we speculate that Jps1 is essential for efficient initiation of exponential growth, eventually by recruiting accessory enzymes like Cts1 to their site of action. The finding that PIP binding and dimerization is conserved in HcJps1 while this version does not complement the *jps1* deletion mutant indicates that further, currently missing species-specific components are involved.

In essence, our study revealed the first detailed insights into the function of Jps1 during unconventional Cts1 secretion and specifically emphasizes the important role of phosphoinositide interaction for these alternative export pathways.

## Experimental procedures

### Accession numbers

The genes and encoding protein sequences are available from Uniprot (https://uniprot.org) with the following accession numbers: *U. maydis* Jps1 (A0A0D1C3B2), *S. reilianum* Jps1 (A0A2N8UGD0), *H. cylindrosporum* Jps1 (A0A0C3CFZ0).

### Molecular cloning and strain generation

*Escherichia coli* Top10 cells (Invitrogen) were used for cloning purposes. Gibson assembly and Golden Gate cloning strategies adapted for efficient generation of plasmids for protein production in *E. coli* and genetic modification of *U. maydis* were employed (40–42). All plasmids were verified by restriction analysis and sequencing. All oligonucleotides used in this study are listed in Table S2.

Plasmids for protein production in *E. coli* were based on pET22b or pEMGB1 (Table S3). The generation of the plasmid pET22b_Jps1_6xHis (pUMa3257) was done using the oligonucleotides oMB945 and oMB946 for amplification of *jps1* from genomic DNA of the sequenced *U. maydis* strain UM521 (Table S3). For generation of the plasmids pEMGB1_Jps1_1-218 (pUX84), pEMGB1_Jps1_1-484 (pUX85), pUMa3257 was used for the amplification of respective *jps1* gene regions. Subsequently, the backbone of pEMGB1_mScarlet (pIL17) (43) was used for *Bsa*I mediated Golden Gate cloning and respective amplified genes were inserted into it. Similarly, genes encoding Jps1 orthologs and Jps1 variants M^1^-M^5^ were cloned into pEMGB1 backbone. *S. reilianum* genomic DNA was isolated from the strain *Sporisorium reilianum* SRZ2 (CBS 131459) (44) and used as template for PCR reactions with the described oligonucleotides to amplify *S. reilianum jps1* gene, which was then further cloned into pEMGB1 backbone to obtain pEMGB1_SrJps1 (pUX91). The gene sequence of *H. cylindrosporum jps1* was dicodon optimized for expression in *U. maydis* (45) and ordered as a gBlock (Integrated DNA Technology, Coralville, Iowa, United States). The optimized sequence was used as a template for PCR reactions with the described oligonucleotides to generate *H. cylindrosporum jps1* gene, which was then further cloned into pEMGB1 to obtain pEMGB1_HcJps1 (pUX122). To generate the Jps1 KR mutants M1-M5, the gene sequence for pUX85 was used as a template for PCR reactions with the described oligonucleotides to amplify the *jps1_1-484* gene with the respective point mutations (Fig. S11). Gene variants were then further cloned into pEMGB1 backbone to obtain pEMGB1_Jps1^1-484^M^1^ (pUX324), pEMGB1_Jps1^1-484^M^2^ (pUX325), pEMGB1_Jps1^1-484^M^3^ (pUX326), Jps1^1-484^M^4^ (pUX185), pEMGB1_Jps1^1-484^M^5^ (pUX186).

Plasmids for genetic modification in *U. maydis* contained a resistance cassette for selection and flanking regions of about 1 kb for homologous recombination, yielding stable strains. For generation of complementation strains, initially the backbone of pPjps1_ Jps1_eGfp_CbxR (pUMa3293) (22) was used to generate pPjps1_Jps1_eGFP_nosT_T2857A_ CbxR (pUX168) based on site-directed mutagenesis to eliminate a EcoRI site. This plasmid was then hydrolyzed with BamHI and EcoRI to be used as a vector backbone for generating plasmids pPjps1_SrJps1_eGfp_CbxR (pUX169), pPjps1_HcJps1_eGfp_CbxR (pUX171), pPjps1_Jps1_1-484_eGfp_CbxR (pUX172), pPjps1_Jps1_1-218_eGfp_CbxR (pUX178), pPJps1_Jps1^1-484^M^5^_eGfp_CbxR (pUX266). The respective *jps1* genes in these plasmids were amplified using Gibson Assembly oligonucleotides as described in the Table S2. Linear constructs obtained by hydrolysis of the respective plasmids were used to transformation *U. maydis* laboratory strain AB33 or derivatives (46). Genetic modifications were verified by Southern blot analysis using the flanking regions as probes. All *U. maydis* strains, genotypes and plasmids used for genetic engineering are listed in Table S4.

### Cultivation

*U. maydis* strains were grown at 28°C in complete medium (CM) supplemented with 1% (w/v) glucose (CM-glc) using baffled flasks and constant shaking at 200 rpm (47). Solid media were supplemented with 2% (w/v) agar. CM-glc agar plates were used for growing the strains on plates.

### Determination of chitinase activity

For determination of extracellular chitinase activity, liquid assays were performed with intact cells according to published protocols (18, 48). An overnight 5 mL CM-glc pre-pre-culture was grown at 28°C, followed by a pre-culture grown over the day. The main culture (10 mL) was then started from this pre-culture to reach an OD_600_ of 1.0 overnight (exception: low density experiments were conducted with cultures grown to OD_600_ 0.15). 2 mL culture was then harvested at 5,000 rpm for 5 min and the pellet was resuspended in 1 mL KHM buffer (110 mM potassium acetate, 20 mM HEPES, 2 mM MgCl_2_). The OD_600_ of this suspension was documented for the analysis of data. 30 µL of each sample was transferred to a black 96-well plate (96 well, PS, F-Bottom, μCLEAR, black, CELLSTAR). MUC, a fluorogenic substrate 4-methylumbelliferyl-β-D-N, N’, N’’-triacetylchitotrioside (MUC, Sigma) (29) was dissolved in DMSO to obtain a stock of 2 mg/mL. The reaction was then initiated by adding 70 µL of MUC working solution containing MUC diluted with KHM buffer (1:10) to 30 µL of the samples. Fluorescence was determined at excitation/emission wavelengths of 360/450 nm at intervals of 5 min for 1 hour at 25°C using the Tecan Infinite M200 plate reader (Männedorf, Switzerland). The gain of the measurement was adjusted for each measurement (gain optimal). Obtained fluorescence values were normalized using OD_600_. For determination of Cts1 activity in the cell extracts, the native cell extracts were prepared as mentioned in the section for Western blot analysis. The total protein concentration of these cell extracts was adjusted to 33 µg/mL using KHM buffer. 30 µL of each extract were then analyzed for Cts1 activity as per the protocol mentioned above.

### Protein production and purification in *Escherichia coli*

*E. coli* BL21 (DE3) (Invitrogen/Life Technologies) and Rosetta(DE3)pLyS cells (Novagen, Merck Millipore) were transformed with plasmid of interest to produce proteins fused with an N-terminal hexa-histidine (His) tag or proteins carrying both GB1 and His-tag. Transformed cells were grown on dYT-agar plates supplemented with 100 µg/mL ampicillin for *E. coli* BL21(DE3) and with 100 µg/mL ampicillin and 34 µg/mL chloramphenicol for Rosetta (DE3)pLyS. Colonies from the plate were used to inoculate a pre-culture in 100 mL dYT medium with appropriate antibiotics grown for 16 h at 37°C under constant shaking at 200 rpm. According to the requirements of the protein of interest, the induction was done either by 1% (w/v) lactose or 0.5 mM isopropyl-β-D-1-thio-galactopyranoside (IPTG, Sigma Aldrich) (**Table 1**). For lactose induction, the main culture was inoculated and subsequently grown at 30 °C for 20 h at 200 rpm. For IPTG induction, a preculture was inoculated and grown at 37°C for 12-16 h which was then used to start a main culture at OD_600_ 0.1, incubated until the OD_600_ reached 0.6. The cultures were then cooled down to 20 °C or incubated at 37 °C and protein production was induced by adding 0.5 mM IPTG. The cells continued to grow for 20 h at 20°C or 3 h at 37 °C and 200 rpm. The cultures were harvested by centrifugation (4,000 g, 15 min, 4°C), resuspended in buffer A (20 mM HEPES pH 8, 20 mM KCl, 40 mM imidazole and 250 mM NaCl) and subsequently disrupted using a microfluidizer (M110-L, Microfluidics). The cell debris was removed by centrifugation (50,000 g, 20 min, 4 °C). The supernatant was loaded onto Ni-NTA FF-HisTrap columns (Cytiva) for affinity purification via the His-tag. The columns were washed with buffer A (10x column volume) and eluted with buffer B (20 mM HEPES pH 8, 20 mM KCl, 250 mM imidazole and 250 mM NaCl). Notably, for 6xHis-Jps1-Gfp a step-wise elution was followed using buffer B with 100, 200, 500 mM imidazole. Eventually, prior to size-exclusion chromatography (SEC), the GB1-tag was cleaved off by adding 0.4 mg TEV protease directly to the eluate and incubating under constant rotation at 20°C for 3 h. Cleaved His-tagged GB1 fragment and remaining TEV were removed via a second Ni-NTA purification after buffer exchange to buffer A using an Amicon Ultra-10K centrifugal filter (Merck Millipore). The tag-free protein was subjected to SEC using a Superdex 200 Increase 26/600 column equilibrated in HEPES buffer (20 mM HEPES pH 7.5, 20 mM KCl and 200 mM NaCl). The peak fractions were analyzed using a standard SDS-PAGE protocol, pooled, and concentrated with Amicon Ultra-10/30/50K centrifugal filters.

**Table 1.** Specific induction conditions for the different recombinant proteins produced in *E. coli*.

| **Protein** | ***E. coli* strain used** | **Induction conditions** | **Buffer used for purification** | **Usage** |
| --- | --- | --- | --- | --- |
| 6xHis-Jps1 | Rosetta(DE3) pLyS | 0.5 mM IPTG, 18°C/18 h | HEPES Buffer | MALS, mass photometry, lipid-binding assays |
| 6xHis-Jps1-Gfp | Rosetta(DE3) pLyS | 0.5 mM IPTG, 37°C/3 h | HEPES Buffer with 5% glycerol | Lipid-binding assays |
| 6xHis-GB1-Jps1<sup>1-484</sup> | Rosetta(DE3) pLyS | 1% lactose (w/v), 30°C/20 h | HEPES Buffer | MALS, mass photometry, lipid-binding assays, SAXS analysis |
| 6xHis-GB1-Jps1<sup>1-218</sup> | Rosetta(DE3) pLyS | 0.5 mM IPTG, 37°C/3 h | HEPES Buffer | MALS, mass photometry, lipid-binding assays |
| 6xHis-GB1-SrJps1 | BL21(DE3) | 1% lactose (w/v), 30°C/20 h | HEPES Buffer | MALS, mass photometry, lipid-binding assays |
| 6xHis-GB1-HcJps1 | BL21(DE3) | 1% lactose (w/v), 30°C/20 h | HEPES Buffer | MALS, mass photometry, lipid-binding assays |
| **6xHis-GB1-Jps1<sup>1-484</sup>M<sup>1</sup>** | Rosetta(DE3) pLyS | 1% lactose (w/v), 30°C/20 h | HEPES Buffer | lipid-binding assays |
| **6xHis-GB1-Jps1<sup>1-484</sup>M<sup>2</sup>** | Rosetta(DE3) pLyS | 0.5 mM IPTG, 37°C/3 h | HEPES Buffer | lipid-binding assays |
| **6xHis-GB1-Jps1<sup>1-484</sup>M<sup>3</sup>** | Rosetta(DE3) pLyS | 1% lactose (w/v), 30°C/20 h | HEPES Buffer | lipid-binding assays |
| **6xHis-GB1-Jps1<sup>1-484</sup>M<sup>4</sup>** | Rosetta(DE3) pLyS | 0.5 mM IPTG, 37°C/3 h | HEPES Buffer | lipid-binding assays |
| **6xHis-GB1-Jps1<sup>1-484</sup>M<sup>5</sup>** | Rosetta(DE3) pLyS | 0.5 mM IPTG, 37°C/3 h | HEPES Buffer | lipid-binding assays |

### Multi-angle light scattering

MALS was performed after SEC purification of the protein of interest. The collected sample was applied onto another SEC column (Superdex 200 GL 10/30; GE Healthcare) which was directly connected to a triple-light scattering detector (miniDAWN TREOS, Wyatt Technology Europe GmbH, Dernbach, Germany) and a differential refractive index detector (OPTILab T-rEX, Wyatt Technology). Astra7 (Wyatt Technology) was used for evaluation of the collected data.

### Mass photometry

Mass photometry experiments were performed using a TwoMP mass photometer (Refeyn Ltd, Oxford, UK). Data acquisition was performed using AcquireMP (Refeyn Ltd. v2.3). Mass photometry movies were recorded at 1 kHz, with exposure times varying between 0.6 and 0.9 ms, adjusted to maximize camera counts while avoiding saturation. Microscope slides (70 x 26 mm) were cleaned for 5 min in 50% (v/v) isopropanol (HPLC grade in Milli-Q H_2_O) and pure Milli-Q H_2_O, followed by drying with a pressurized air stream. Silicon gaskets to hold the sample drops were cleaned in the same manner and fixed to clean glass slides immediately prior to measurement. The instrument was calibrated using the NativeMark Protein Standard (Thermo Fisher Scientific) immediately prior to measurements. The concentration during measurement of UmJps1 full length and variants as well as Jps1 orthologs was typically 10 nM. Each protein was measured in a new gasket well (i.e., each well was used once). To find focus, 18 μL of fresh HEPES buffer adjusted to room temperature was pipetted into a well, the focal position was identified and locked using the autofocus function of the instrument. For each acquisition, 2 μL of diluted protein was added to the well and thoroughly mixed. For each sample, three individual measurements were performed. The data were analyzed using the DiscoverMP software.

### Hydrogen/deuterium exchange mass spectrometry (HDX-MS)

HDX-MS samples were prepared by a two-arm autosampler (LEAP Technologies). Reactions were initiated by adding 58.5 µL of buffer prepared with D_2_O (20 mM HEPES-Na pH 7.5, 20 mM KCl, 20 mM MgCl_2_, 200 mM NaCl), to 6.5 µL of Jps1 solution (50 µM). After incubation at 25 °C for 10, 95, 1,000 or 10,000 s, HDX was quenched by transferring 55 µL of the reaction to another well containing 55 µL of 400 mM KH_2_PO_4_/H_3_PO_4_, 2 M guanidine-HCl (pH 2.2) kept at 1 °C. After mixing, 100 μL of the quenched reaction were injected into an ACQUITY UPLC M-Class system with HDX Technology (49). Non-deuterated samples were generated similarly with an H_2_O-based buffer. The samples were washed from a 50-μL injection loop with water + 0.1% (v/v) formic acid (100 μL/min) and passed through a cartridge (2 mm x 2 cm, kept at 12 °C) filled with porcine pepsin immobilized to bead material. The resulting peptic peptides were collected on a trap column (2 mm x 2 cm; 0.5 °C) filled with POROS 20 R2 (ThermoFisher Scientific). After 3 minutes, the column was placed in line with an ACQUITY UPLC BEH C18 1.7 μm 1.0 x 100 mm column (Waters) temperated at 0.5 °C and the peptides eluted with a gradient of H_2_O + 0.1% (v/v) formic acid (eluent A) and acetonitrile + 0.1% (v/v) formic acid (eluent B) at 60 μL/min, as follows: 0-9 min/95-55% A, 9-10 min/55-15% A, 10-10.1 min/15-5% A, 10.1-11/5% A. Peptides were guided to a Synapt G2-Si mass spectrometer (Waters) and ionized by electrospray ionization (capillary temperature: 250 °C; spray voltage: 3.0 kV). Mass spectra were acquired with MassLynX MS 4.1 (Waters) over 50 to 2,000 *m/z* in enhanced high-definition MS (HDMS^E^) (50, 51) or high-definition MS (HDMS) mode for non-deuterated and deuterated samples, respectively. Lock-mass correction was conducted with [Glu1]-fibrinopeptide B standard (Waters). During peptide separation on the ACQUITY UPLC BEH C18 column, the pepsin column was washed 3 times with 80 μL of 0.5 M guanidine-HCl in 4% (v/v) acetonitrile. Blank runs (double-distilled H_2_O) were performed between each sample. All measurements were performed in duplicate (separate HDX reactions).

Peptides were identified with ProteinLynx Global SERVER 3.0.1 (PLGS, Waters) from the non-deuterated samples acquired with HDMS^E^ employing low-energy, elevated-energy and intensity thresholds of 300, 100 and 1,000 counts, respectively, and matched using a database containing the amino acid sequences of Jps1, porcine pepsin, and their reversed sequences (peptide tolerance = automatic; fragment tolerance = automatic; min fragment ion matches per peptide = 1; min fragment ion matches per protein = 7; min peptide matches per protein = 3; maximum hits to return = 20; maximum protein mass = 250,000; primary digest reagent = non-specific; missed cleavages = 0; false discovery rate = 100). For quantification of deuterium incorporation with DynamX 3.0 (Waters), peptides had to fulfil the following criteria: identification in both non-deuterated samples; minimum intensity of 20,000 counts; maximum length of 40 residues; minimum number of two products; maximum mass error of 25 ppm; retention time tolerance of 0.5 minutes. All spectra were manually inspected and omitted, if necessary, e.g., for low signal-to-noise ratios or overlapping peptides prohibiting correct assignment of the isotopic clusters.

Residue-specific deuterium uptake from peptides identified in the HDX-MS experiments was calculated with the software DynamX 3.0 (Waters). In the case that any residue is covered by a single peptide, the residue-specific deuterium uptake is equal to that of the whole peptide. In the case of overlapping peptides for any given residue, the residue-specific deuterium uptake is determined by the shortest peptide covering that residue. Where multiple peptides are of the shortest length, the peptide with the residue closest to the peptide C-terminus is utilized. Assignment of residues exhibiting no higher-order structure (disordered) was based on two criteria, i.e., a residue-specific deuterium uptake of >50% after 10 s of HDX and no increment in HDX >5% in between consecutive HDX time-points (52). Raw data of deuterium uptake by peptides and residue specific HDX are provided in **Supplementary Dataset 1** (53).

### Small-angle X-ray scattering

SEC-SAXS data were collected on the P12 beamline (PETRA III, DESY Hamburg (54)). The sample to detector distance of the P12 beamline for was 3.00 m, resulting in an achievable q-range of 0.03 – 4.4 nm^-1^. The measurement was performed at 10°C with a protein concentration of 0.6 mg/mL of purified Jps1^1-484^. The sample was measured in batch mode and injected via autosampler. We collected 40 frames with an exposer time of 0.095 sec/frame. Data were collected on absolute scale intensity against water.

All used programs for data processing were part of the ATSAS Software package (Version 3.0.5) (55). Primary data reduction was initially performed with the SASFLOW (56) pipeline and checked with the programs PRIMUS (57). The Guinier approximation (58) was used to determine the forward scattering *I*(*0*) and the radius of gyration (*R_g_*). The pair-distribution function *p(r)* was created with the program GNOM (59) and determined the maximum particle dimension (*D_max_*). The rigid body modelling of the Jps1^1-484^ dimer was done with CORAL (60). We used an AlphaFold2 (24, 61) prediction as template, where the flexible loop regions (aa 357-447) as well as the N-terminal residues (aa 1-17, 75-172) of the dimer were remodeled via CORAL (60).

### PIP overlay assays

The lipid binding analysis of Jps1 was performed using PIP Lipid Strips (Echelon Biosciences Inc.). Prior to their use, the lipid strips were blocked for one hour in 1x TBS buffer with 0.05 % Tween 20 (TBS-T) supplemented with 3 % (w/v) BSA. 5 μg of purified Jps1 protein was incubated with the pre-blocked strips in TBS-T buffer supplemented with 3% BSA in the dark (total volume 15 mL). Next, unbound protein was removed in three 10 min washing steps with TBS-T. An anti-His-primary antibody (Sigma, 1:2,000) was incubated with the protein-treated lipid strips to detect Jps1 bound via its C-terminal His-tag. The unbound antibody was then removed by subsequent washing steps after incubation. AceGlow Western Blot detection solution was used to detect chemiluminescent signals on the lipid membrane. 0.5 µg PI(4,5)P_2_-grip protein (GST-tagged PLC-δ1 PH domain protein, Echelon) was used as positive control. Here, a primary anti-GST (Sigma, 1:3,000) was used to detect bound PI(4,5)P_2_-grip protein.

### Liposome binding assays

DOPC (1,2-di-(9Z-octadecenoyl)-sn-glycero-3-phosphocholine) (catalog no. 850375C-25MG), cholesterol (3β-hydroxy-5-cholesten, 5-cholesten-3β-ol) (catalog no. C8667-1G) and PI(4,5)P_2_ (L-α-phosphatidylinositol-4,5-bisphosphate (Brain, Porcine) ammonium salt) (catalog no. 840046P-1MG) were purchased from Sigma Aldrich/Merck (manufactured by Avanti Polar Lipids, Inc.). Lipids solved in chloroform were mixed in desired concentrations and ratios, and chloroform was evaporated under vacuum conditions at 40°C using a rotary evaporator (IKA). The dried lipid film was then resuspended in 20 mM HEPES pH 7.5 buffer supplemented with 20 mM KCl and 200 mM NaCl to achieve the lipid concentration of 5 mM. In order to obtain large unilamellar vesicles, the crude liposomes were manually extruded through porous polycarbonate membranes (Nucleopore, Whatman) using the Mini-Extruder set (Avanti Polar Lipids, Inc.). The membrane pore size was decreased stepwise from 200 nm to 50 nm. The control liposomes were prepared by using 100 mol % DOPC and DOPC / cholesterol in the molar ratio 80:20. Liposomes containing phosphoinositides were prepared by mixing DOPC:cholesterol:PIP and DOPC:PIP in the molar ratios 80:15:5 and 95:5, respectively.

For liposome binding assays, the protein of interest was mixed with liposomes in a protein: lipid molar ratio of 1:5,000 in a final volume of 100 µL made up with HEPES buffer (20 mM HEPES pH 7.5, 20 mM KCl and 200 mM NaCl). This suspension was incubated at room temperature for 15 min to allow the protein-liposome interaction. After incubation, the suspension was ultra-centrifuged at 52,000 rpm for 45 min. The pellet and supernatant fractions were then carefully separated and the pellet was resuspended in 100 µL HEPES buffer. The pellet and supernatant fractions were then precipitated by addition of 20% (w/v) TCA and further analyzed using standard SDS-PAGE protocol followed by Coomassie staining.

### Binding studies using giant unilamellar vesicles

All lipids were purchased from Cayman Chemicals or Avanti Polar Lipids, and stock solutions (0.2 mg/mL, except for cholesterol 10 mg/mL) were prepared with chloroform. GUVs were produced using PVA-assisted swelling (62). 100 μL of 5% (w/w) polyvinyl alcohol (PVA) were spread equally on the front of an object slide within a defined area. For drying, the PVA coated slide was placed on a thermoblock heated to 50°C. Lipids were mixed in the following molar ratios – DOPC (74.75 mol%), cholesterol (20 mol%), PIP (5 mol%), Texas Red (0.25 mol%). 10 μL of the lipid mixes were spread equally onto the PVA coated slides and another 10 µL was added upon the subsequent drying of the initial layer. Marked areas of PVA + dried lipids were surrounded by Vitrex on the slides in a “U”-shape. The second cleaned object slide was squeezed onto the Vitrex, generating a sealed chamber. Approximately, 500 μL of 10% (w/v) sucrose solution were filled carefully into the chamber. The sucrose-filled chamber was sealed at the top with another layer of Vitrex. Swelling was performed at room temperature (∼25°C) for one hour. GUVs were harvested from the incubation chamber with microcapillary-tips (VWR) transferred into fresh reaction tubes and analyzed by microscopy.

For binding experiments, 10 μL of GUVs were used and mixed with different amounts of protein. 30-300 μg of 1 mg/mL purified Jps1G were used for the GUV binding studies. Total protein concentration in the well was adjusted with 1x TBS buffer (max. volume of plate wells: 50 μL). To immobilize GUVs and to avoid interaction of Jps1 with the negatively charged surface of the wells, Ibidi® plate wells were coated with 30 μL of β-casein solution for five minutes at room temperature. β-casein was discarded and wells were washed with 50 μL of 1xTBS-buffer three-times. These plates either coated with casein or uncoated were used for inverted laser scanning confocal microscopy with an AiryScan module (Zeiss).

### Surface plasmon resonance (SPR)

The protein: lipid binding experiments were performed in the two-channel instrument 2SPR (AMETEK Reichert Inc., Depew, NY, USA) using SPR sensor chip LP (coating -lipophilic groups covalently bound to a 2D carboxymethyldextran surface) used for the capture of lipid membrane vesicles (XanTec bioanalytics GmbH, Düsseldorf, Germany). DOPC and DOPC, PI(4,5)P_2_ liposomes were extruded respectively as mentioned before to the diameter of 50 nm. Initially, the chip surface was equilibrated with SPR running buffer (20 mM HEPES pH 7.5, 20 mM KCl and 200 mM NaCl) for 25 min. The chip was then cleaned simultaneously with 20 mM CHAPS (2 injections for 30s each), 50 mM NaOH (2 injections for 30 s each), 2:3 isopropanol: 50 mM NaOH (2 injections for 1 min each) at a flow rate of 25 µL*min^-1^. The liposomes were immobilized at a flow rate of 10 µL*min^-1^ for 10 min to achieve 8,000 to 10,000 response units (RU). DOPC-only liposomes (1 mM) lacking the protein interaction were immobilized first in both the reference and test channel. The switch to the test channel was then made and the immobilized liposomes were removed completely using 2:3 isopropanol: 50 mM NaOH (1 injection for 1 min) until the response comes back to the baseline. After this, the DOPC, PI(4,5)P_2_ (1 mM) liposomes were immobilized through the test channel. The switch to both channels was then made and 1 injection of 50 mM NaOH was done to remove any loosely bound liposomes. Further, BSA (0.5 mg/mL) was injected for 1 min to check the specificity of the liposomes. The flow rate was then switched to 25 µL*min^-1^ and the desired concentration of the protein of interest (200 µL) was injected. After each measurement, the chip was regenerated using the cleaning procedure mentioned above and further desired concentrations of protein were tested following the same procedure. Data was analysed using TraceDrawer (Version 1.9.2) software program (Ridgeview Instruments AB, Uppsala, Sweden). The data set was fitted by non-linear regression analysis of response units using the steady state affinity model: Y = B_max_ * c/ (c + K_D_), where K_D_ is the dissociation constant, B_max_ is the maximum response unit, c is the concentration of HcJps1.

### Western blot analysis

To evaluate the production of Jps1 variants in cell extracts, 50 mL cultures were grown to an OD_600_ of 1.0 and harvested at 5000 rpm for 5 min at 4°C. For intracellular Cts1 activity assays, the cell pellets were then resuspended in 1 mL of native extraction buffer containing 1X PBS pH 7.2, 100 µL 0.1 M PMSF (Sigma/Aldrich), 100 µL Protease inhibitor cocktail (Roche), 50 µL 0.5 mM benzamidine (Sigma Aldrich) and 400 µL of glass beads were added to each tube. The cells were then disrupted using Retsch Mill at 30 Hz for 15 min. After the cell disruption, the cell extracts were then centrifuged at 13,000 rpm for 30 min at 4°C to settle down the cell debris. The cell extracts were then transferred to fresh reaction tubes and used for further analysis. Alternatively, after harvesting the cultures, the cell pellets were resuspended in 2 mL PBS pH 7.2 and transferred to fresh centrifuge tubes. The cells were the harvested at 5,000 rpm for 5 min and the supernatant was removed completely. The resulting cell pellets were flash-frozen in liquid nitrogen, Sample tubes were placed on 24 well TissueLyser adapter

Qiagen 69982) and soaked in liquid nitrogen for 1 min, followed by further addition of 5 mm stainless steel bead to each sample tube. The cells were disrupted using Retsch Mill at 30 Hz for 1 min X 3 cycles. After this, the dry homogenized powder of cells was resuspended in 1 mL urea buffer (8 M urea, 50 mM Tris/HCl pH 8.0 containing one tablet of ‘complete protease inhibitor” per 25 mL, 1 mM DTT, 0.5 mM benzamidine) and further centrifuged at 13,000 rpm for 10 minutes at 4°C. The supernatant was then used for subsequent analysis. Protein concentrations were determined by Bradford assay (BioRad, Hercules, CA, United States). 50 μg of total protein was analysed using standard SDS-PAGE protocol and transferred to a PVDF membrane (Amersham, Hybond -P), activated in 100% methanol using semi-dry Western blotting. Gfp-tagged protein of interest was detected using a primary mouse anti-Gfp (1:3,000, Millipore/Sigma). An anti-mouse IgG-horseradish peroxidase (HRP) conjugate (1:3,000 Promega) was used as secondary antibody. HRP activity was then detected using the Amersham ECL Primer Western Blotting Detection Reagent (Cytiva) and a LAS4000 chemiluminescence imager (GE Healthcare).

### Microscopy and staining procedures

The microscopic analysis of an overnight grown culture (CM-glc) at OD_600_ of 0.5 was performed using wide-field fluorescence microscope Zeiss Axio Imager M1 equipped with a Spot Pursuit CCD camera (Diagnostic Instruments, Sterling Heights, MI) and objective lenses Plan Neofluar (63x, NA 1.25) and Plan Neofluar (100x, NA 1.40). The fluorescent proteins were excited with a HXP metal halide lamp (LEj, Jena) in combination with filter sets for Gfp (ET470/40BP, ET495LP, ET525/50BP), mCherry (ET560/40BP, ET585LP, ET630/75BP Chroma, Bellow Falls, VT), and DAPI (HC387/11BP, BS409LP, HC 447/60BP; AHF Analysentechnik, Tübingen, Germany). The system was operated by MetaMorph, version 7 (Molecular Devices, Sunnyvale, CA, United States). Image processing including the adjustments of brightness and contrast were conducted by ImageJ software version 1.54g. To visualize the fungal septa yeast cells were stained by addition of calcofluor white to the culture (1 µg/ml) before microscopy. Airyscan microscopy was applied using a Zeiss inverted LSM880 airyscan microscope system (Zeiss Microscopy GmbH, Oberkochen, Germany), equipped with a Plan-Apochromat 40x/1.2 water objective lens. Images were acquired using fast airyscan mode of the airyscan1 module. The general acquisition parameters were set as the following. 488 nm was used at 10% intensity as an excitation laserline for eGFP/Jps1-eGFP with a BP 465-505 + 525+555 nm detection filter. 561 nm was used at 1% intensity as an excitation laserline for the the TexasRed labeled GUVs with a BP 570-620 + LP 645 detection filter. The airyscan detector gain was set to 900 for eGFP/Jps1-eGFP and to 750 for TexasRed labeled GUVs. The scans were performed in unidirectional frame-sequential mode at a pixel dwell time of 0.98-1.96 µsec/pixel and a pixelsize of 99 nm. The final data were calculated using the Zeiss built in airyscan module and automatic airyscan strength parameter were varying between 2.7-2.9 for eGFP/Jps1-eGFP and between 3.0-3.3 for TexasRed labeled GUVs.

### Bioinformatics

For structure prediction of UmJps1 and orthologs, the respective sequences of the full-length proteins were downloaded from the UniProt database and AlphaFold2 v2.3 was used in Multimer mode using default settings (24).

## Data Availability

HDX-MS raw data are provided in the Supporting Information as Supplementary Dataset 1. We uploaded the SAXS data to the Small Angle Scattering Biological Data Bank (SASBDB) (63), with the accession codes SASDT97.

## Supporting information

This article contains supporting information.

## Acknowledgements

We are grateful to Bettina Axler for excellent technical support and Michael Feldbrügge for critical reading of the manuscript. We acknowledge the DESY (Hamburg, Germany), a member of the Helmholtz Association HGF, for the provision of experimental facilities. Parts of this research were carried out at PETRA III and we would like to thank Dmytro Soloviov (EMBL Hamburg) for the assistance in using beamline P12. We are grateful to Sebastian Joecks and Refeyn Ltd. for access to and assistance with the TwoMP mass photometer. We thank Eymen Hachani from the Institute of Biochemistry I at Heinrich-Heine University for conducting MALS analyses.

## Funding and additional information

This work was funded by the German Research Foundation (DFG) within the Collaborative Research Center 1208 (project ID 267205415, sub-project A09/Schipper) and 1535 (project ID 458090666, sub-project Z01 to S.S. and J.R.). The Center for Structural studies is funded by the DFG (Grant number 417919780 and INST 208/761-1 FUGG to S.S.). We thank the German Research Foundation (DFG) for support through the Core Facility for Interactions, Dynamics and Macromolecular Assembly (project 324652314). We would like to acknowledge the Center for Advanced Imaging (CAi) at Heinrich-Heine-University Düsseldorf for providing access to the Zeiss LSM880 Airyscan system (Ref.No. DFG-INST 208/746-1 FUGG).

