## Supplementary File for "The fungal protein Jps1 facilitates unconventional protein secretion through a direct phosphoinositide interaction"

### Running title

Insights into the lipid interaction mechanism of Jps1

### Keywords

cytokinesis, unconventional secretion, protein-lipid interaction, fungi, liposome, membrane binding

**Table S1.** Overall SAXS data from Jps1$^{1-484}$.

| | |
|---|---|
| **Data collection parameters** | |
| SAXS Device | P12, PETRA III, DESY Hamburg (1) |
| Detector | PILATUS 6 M (423.6 x 434.6 $mm^2$) |
| Detector distance (m) | 3.0 |
| Beam size | 120 μm x 200 μm |
| Wavelength (nm) | 0.124 |
| Sample environment | Quartz glass capillary, 1 mm ø |
| Absolute scaling method | Comparison with scattering from pure $H_2O$ |
| Normalization | To transmitted intensity by beam-stop counter |
| Scattering intensity scale | Absolute scale, $cm^{-1}$ |
| *s* range ($nm^{-1}$)‡ | 0.03 – 4.4 |
| **Sample** | |
| Organism | *Ustilago maydis* (strain 521 / FGSC 9021) |
| UniProt ID (range) | A0A0D1C3B2 (1-484) |
| Mode of measurement | batch |
| Temperature (°C) | 10 |
| Exposure time (# frames) | 0.095 (40) |
| Protein buffer | 20 mM HEPES pH 7.5, 20 mM KCl, 200 mM NaCl |
| Protein concentration (mg/ml) | 0.6 |
| **Structural parameters** | |
| *Guinier Analysis (PRIMUS)* | |
| $I(0) \pm \sigma$ ($cm^{-1}$) | 0.093 ± 0.00025 |
| $R_g \pm \sigma$ (nm) | 4.05 ± 0.02 |
| *s-range* ($nm^{-1}$) | 0.066 – 0.320 |
| *min < sRg < max limit* | 0.268 – 1.294 |
| Data point range | 1 - 89 |
| Linear fit assessment ($R^2$) | 0.992 |
| *PDDF/P(r) Analysis (GNOM)* | |
| $I(0) \pm \sigma$ ($cm^{-1}$) | 0.094 ± 0.00027 |
| $R_g \pm \sigma$ (nm) | 4.20 ± 0.02 |
| $D_{max}$ (nm) | 14.65 |
| Porod volume ($nm^3$) | 253.61 |
| *s-range* ($nm^{-1}$) | 0.066 – 4.087 |
| $\chi^2$ / CorMap P-value | 0.950 / 0.746 |
| **Molecular mass (kDa)** | |
| From *I*(0) | 119.86 |
| From Vc (2) | 110.03 |
| Bayesian Inference (3) | 130.88 |
| GNNOM (4) | 131.10 |
| From sequence | 60.80 (monomer)<br>121.60 (dimer) |
| **Rigid body modeling** | |
| CORAL | |
| Symmetry | P1 |
| *s*-range for fit ($nm^{-1}$) | 0.066 – 4.087 |
| $\chi^2$, CorMap *P*-value | 1.06 / 0.0026 |
| **SASBDB accession codes** (5) | |
| **Software** | |
| ATSAS Software Version (6) | 3.0.5 |
| Primary data reduction | SASFLOW [3] / PRIMUS (7) |
| Data processing | GNOM (8) |
| *Rigid body* modelling | CORAL (9) |
| Model visualization | PyMOL (10) |

‡$s = 4\pi\sin(\theta)/\lambda$, $2\theta$ – scattering angle

**Table S2.** Oligonucleotides used in this study.

| Number | Sequence (5´ to 3`) | Reference |
|---|---|---|
| oCD807 | AGGAGGGTCTCCTCGAGAGCCGAACTACTCGGAGCATCCCA | This work |
| oEF319 | GGTCTCCCATGGGCCCAGGCATCTCCAAGAAGCCTTCTT | This work |
| oEF320 | GGTCTCCTCGAGATCGTCTTGCAGAGCGCCGG | This work |
| oEF333 | GGTCTCCCATGGGCCCAGGCATCTCCAAGAAGCCCTC | This work |
| oEF334 | GGTCTCCTCGAGGGATTCGGCGTCGATGGGCG | This work |
| oEF483 | GGTCTCCTCGAGCCATCGAAAGATGCTCTTTTTTTTCTCC | This work |
| oEF517 | GGTCTCCTCGAGCCAACGAAAAATGGACTTC | This work |
| oEF979 | CCACGACTTGCAAATCGGATCCATGCCAGGCATCTCCAAGAAGCCTTC | This work |
| oEF980 | CCTCGCCCTTGCTCACCATGAATTCAGCCGAACTACTCGGAGCATCCC | This work |
| oEF969 | CCACGACTTGCAAATCGGATCCATGCCAGGCATCTCCAAGAAGCCCTCG | This work |
| oEF970 | CCTCGCCCTTGCTCACCATGAATTCGGATTCGGCGTCGATGGGCGTTCG | This work |
| oEF987 | GATAAGCTGTCAAACATGAGAATACATCGATGATATCAGATCTGC | This work |
| oEF988 | GCAGATCTGATATCATCGATGTATTCTCATGTTTGACAGCTTATC | This work |
| oEF989 | CCACGACTTGCAAATCGGATCCATGGACGATGGCCTGGTCCGCATG | This work |
| oEF990 | CCTCGCCCTTGCTCACCATGAATTCCCAACGAAAAATGGACTTC | This work |
| oEF997 | CCTCGCCCTTGCTCACCATGAATTCATCGTCTTGCAGAGCGCCGG | This work |
| oMB945 | CATATGCCAGGCATCTCCAAGAAGCC | This work |
| oMB946 | CTCGAGGGATTCCGCATCGATTGGGGTTTGG | This work |
| oXA21 | GGTCTCCAAGCCAATGCGGCTGCGGCTTCGTTCGCTCCTTCTGGC | This work |
| oXA22 | GGTCTCGGCTTGCGCCGCTGCCTCCACTGCTGCCGCCTGAGC | This work |
| oXA28 | GGTCTCCGCCGAAGAAGAAAGCTGGACAGGAGTGGG | This work |
| oXA29 | GGTCTCGCGGCCGCCGCCGCGACGACGGTGCCAGAAGGAGCG | This work |
| oXA241 | GGTCTCCGGTGGCGCTGGCGGAGAACCCTCATGTCTCGG | This work |

| | | |
|---|---|---|
| oXA242 | GGTCTCCCACCGTCGCCGCGTGAGGCGAGACGTGAGG ACTGCC | This work |
| oXA243 | GGTCTCGTCGGCGATCTTTACCGGTGGTCATGGATACC CCGGTGTTCC | This work |
| oXA244 | GGTCTCGCCGACGCAGGAGAGAGGAAAGCCGAG | This work |
| oXA245 | GGTCTCGCTCGCGGCTCAGGCGGCAGCAGTGG | This work |
| oXA246 | GGTCTCGCGAGCGCATCGCAGTTCATGATGCTCG | This work |

**Table S3.** Plasmids used in this study. For detailed description see main text.

| **Plasmid (collection no.)** | **Usage** | **Reference** |
|---|---|---|
| pET22b | Protein overexpression with N-terminal 6His-tag | Novagen |
| pEMGB1 (pIL17) | Protein overexpression with N-terminal GB1-tag | (11, 12) |
| pET22b_Jps1_6xHis (pUMa3257) | Protein expression of 6xHis-Jps1 full-length | This work |
| pET15b_6xHis_Jps1_Gfp (pUMa4285) | Protein expression of 6xHis-Jps1Gfp full-length | This work |
| pPjps1_Jps1_eGfp_CbxR (pUMa3293) | Generation of plasmid pUX168 (via site directed mutagenesis) | This work |
| pEMGB1_Jps1_1-218 (pUX84) | Protein expression of GB1-Jps1_1-218 | This work |
| pEMGB1_Jps1_1-484 (pUX85) | Protein expression of GB1-Jps1_1-484 | This work |
| pEMGB1_SrJps1 (pUX91) | Protein expression of GB1-SrJps1 full-length | This work |
| pEMGB1_HcJps1 (pUX122) | Protein expression of GB1-HcJps1 full-length | This work |
| pEMGB1_Jps1$^{1\text{-}484}$M$^{1}$ (pUX324) | Protein expression of GB1-Jps1M1 | This work |
| pEMGB1_Jps1$^{1\text{-}484}$M$^{2}$ (pUX325) | Protein expression of GB1-Jps1M2 | This work |
| pEMGB1_Jps1$^{1\text{-}484}$M$^{3}$ (pUX326) | Protein expression of GB1-Jps1M3 | This work |
| pEMGB1_Jps1$^{1\text{-}484}$M$^{4}$ (pUX185) | Protein expression of GB1-Jps1M4 | This work |
| pEMGB1_Jps1$^{1\text{-}484}$M$^{5}$ (pUX186) | Protein expression of GB1-Jps1M5 | This work |
| pPjps1_eGfp_nosT_T2857A_CbxR (pUX168) | Backbone vector for generation of pUX169, pUX170, pUX171, pUx172, pUX178, pUX266 | This work |
| pPjps1_SrJps1_eGfp_CbxR (pUX169) | Generation of complementation strain for SrJps1-eGfp (UX140) | This work |
| pPjps1_HcJps1_eGfp_CbxR (pUX171) | Generation of complementation strain for HcJps1-eGfp (UX141) | This work |

| | | |
|---|---|---|
| pPjps1_Jps1_1-484_eGfp_CbxR (pUX172) | Generation of complementation strain for Jps11-484-eGfp (UX130) | This work |
| pPjps1_Jps1_1-218_eGfp_CbxR (pUX178) | Generation of complementation strain for Jps11-218-eGfp (UX134) | This work |
| pPjps1_Jps1$^{1-484}$M$^{5}$_eGfp_CbxR (pUX266) | Generation of complementation strain for Jps1M5-eGfp (UX207) | This work |

**Table S4.** Strains used in this study.

| **Strain** | **Internal strain collection number** | **Parental strain** | **Vector** | **Genotype** | **Reference** |
|---|---|---|---|---|---|
| ***U. maydis*** | | | | | |
| AB33 | UMa133 | | - | *a2 Pnar: bW2bE1 PhleoR* | (13) |
| UM521 | UMa54 | | | *a1b1* | (14) |
| AB33 jps1Δ | UMa2092 | UMa133 | | *a2 Pnar: bW2bE1 PhleoR umag_03776Δ_HygR* | (15) |
| AB33 Jps1G | UMa1847 | UMa133 | - | *a2 Pnar: bW2bE1 PhleoR umag03776- egfp_HygR* | (15) |
| AB33 jps1Δ/ Jps1$^{1-484}$G | Ux130 | UMa2092 | pUX172 | *a2 Pnar: bW2bE1 PhleoR umag_03776Δ_HygR_ip$^{r}$(pJps1_Jps1_1-484_eGfp_nosT) ip$^{s}$ CbxR* | This work |
| AB33 jps1Δ/ Jps1$^{1-218}$G | UX134 | UMa2092 | pUX178 | *a2 Pnar: bW2bE1 PhleoR umag_03776Δ_HygR_ip$^{r}$(pJps1_Jps1_1-218_eGfp_nosT) ip$^{s}$ CbxR* | This work |
| AB33 jps1Δ/ SrJps1G | Ux140 | UMa2092 | pUX169 | *a2 Pnar: bW2bE1 PhleoR umag_03776Δ_HygR_ip$^{r}$(pJps1_SrJps1_eGfp_nosT) ip$^{s}$ CbxR* | This work |
| AB33 jps1Δ/ HcJps1G | Ux141 | UMa2092 | pUX171 | *a2 Pnar: bW2bE1 PhleoR umag_03776Δ_HygR_ip$^{r}$(pJps1_HcJps1_eGfp_nosT) ip$^{s}$ CbxR* | This work |
| AB33 jps1Δ/ Jps1$^{1-484}$M$^{5}$G | UX207 | UMa2092 | pUX266 | *a2 Pnar: bW2bE1 PhleoR umag_03776Δ_HygR_ip$^{r}$(pJps1_Jps1$^{1-484}$M$^{5}$_eGfp_nosT) ip$^{s}$ CbxR* | This work |
| **Others** | | | | | |
| *Sporisorium reilianum* SRZ2 | UMa695 (RK106) | | - | *a2b2* | (16) |

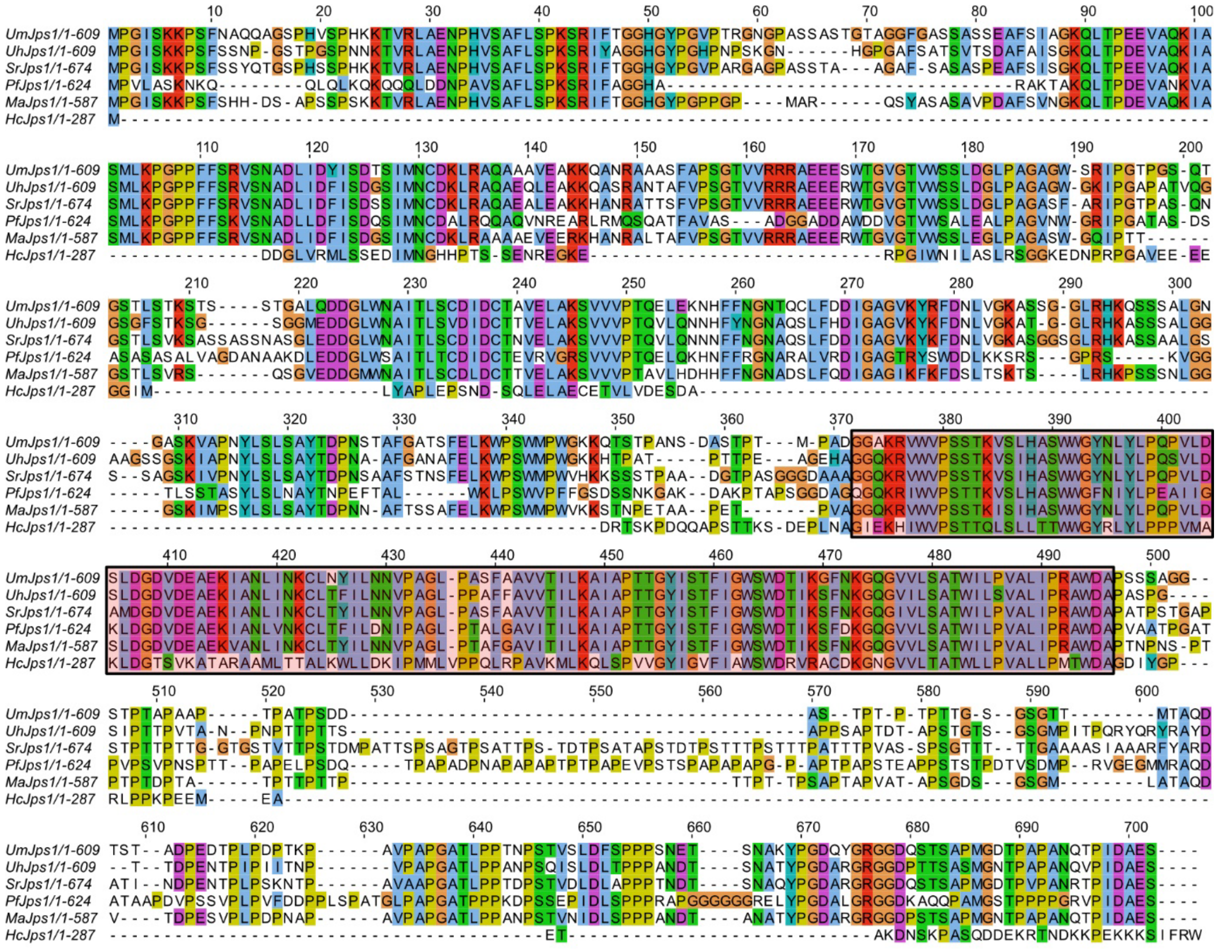

**Supplementary Figure 1. Amino acid alignment of Jps1 and selected homologs.** Homologous sequences were identified by BlastP and examples ranging from close relatives (smut fungi) to more distantly related Basidiomycetes were selected. The boxed region resembles the highly conserved core domain. The alignment was generated using the Clustal Omega webserver (17) and visualized using JalView (18). *Um, Ustilago maydis; Uh, Ustilago hordei; Sr, Sporisorium reilianum; Pf, Pseudozyma flocculosa; Ma,Moesziomyces antarcticus; Hc, Hebeloma cylindrosporium*. Numbers indicate protein lengths (amino acids).

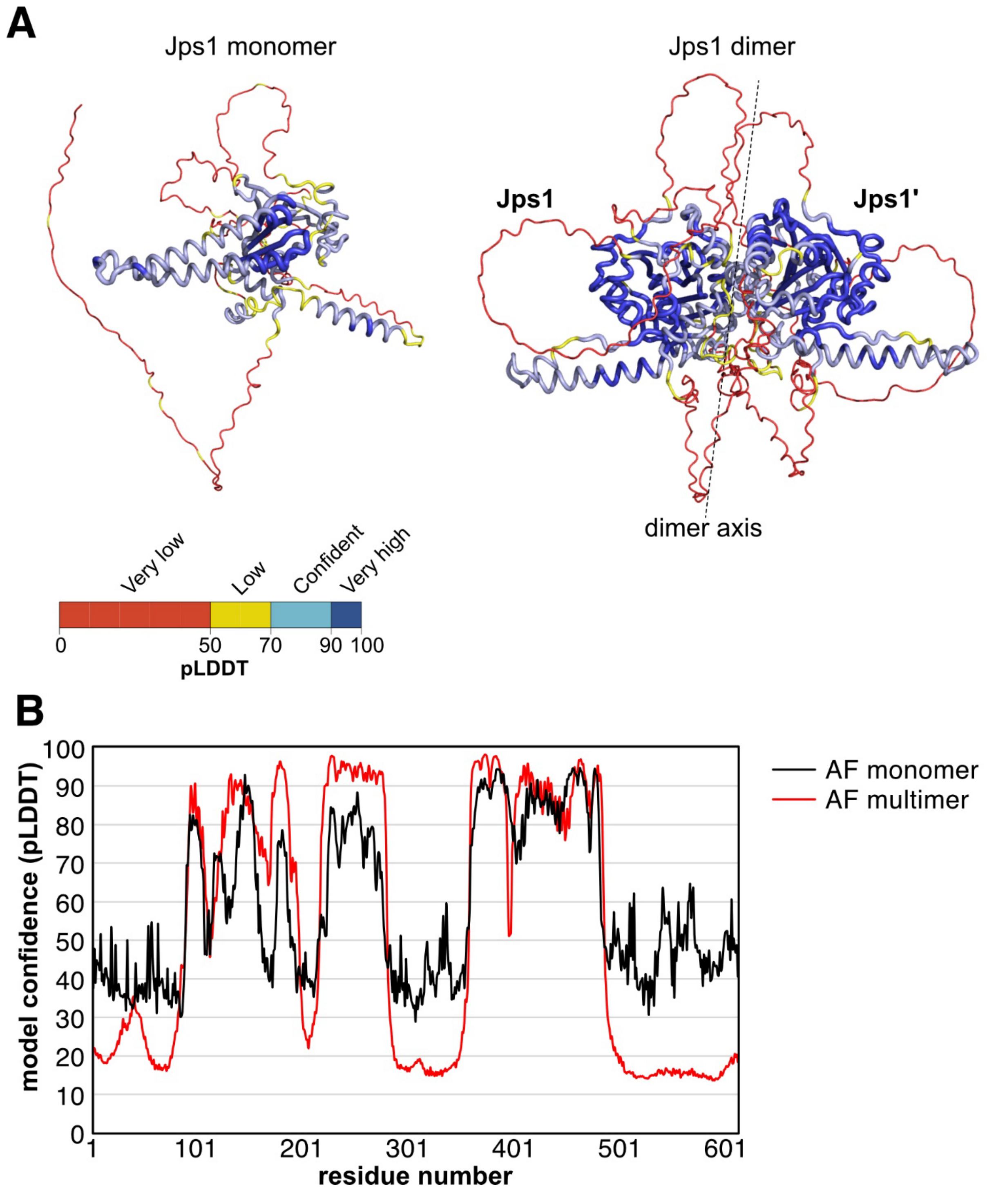

**Supplementary Figure 2. Structure prediction of Jps1 using AlphaFold2. A.** Structural models of the Jps1 monomer (left) and dimer (right) colored according to pLDDT values predicted with AlphaFold2 (19). **B.** Model confidence (pLDDT) score of a Jps1 monomer (black) and dimer (red). The pLDDT is plotted per residue. Notably, the pLDDT in the conserved regions improved from around 80 to >95, while it strongly dropped in the less conserved regions suggesting intrinsically disordered regions.

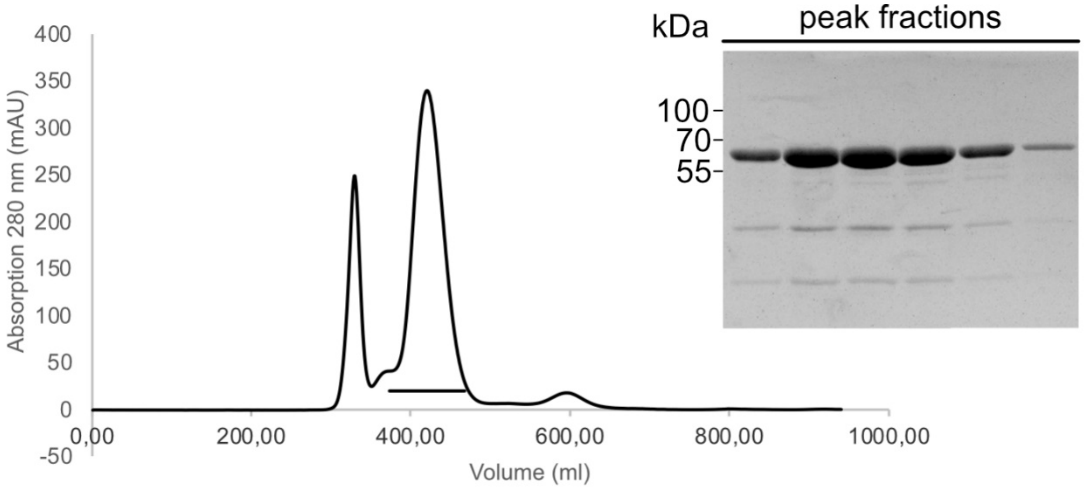

**Supplementary Figure 3. Purification of Jps1.** Shown are a chromatogram of full-length Jps1 from the size-exclusion chromatography and an SDS-PAGE of the peak fractions marked with a black line.

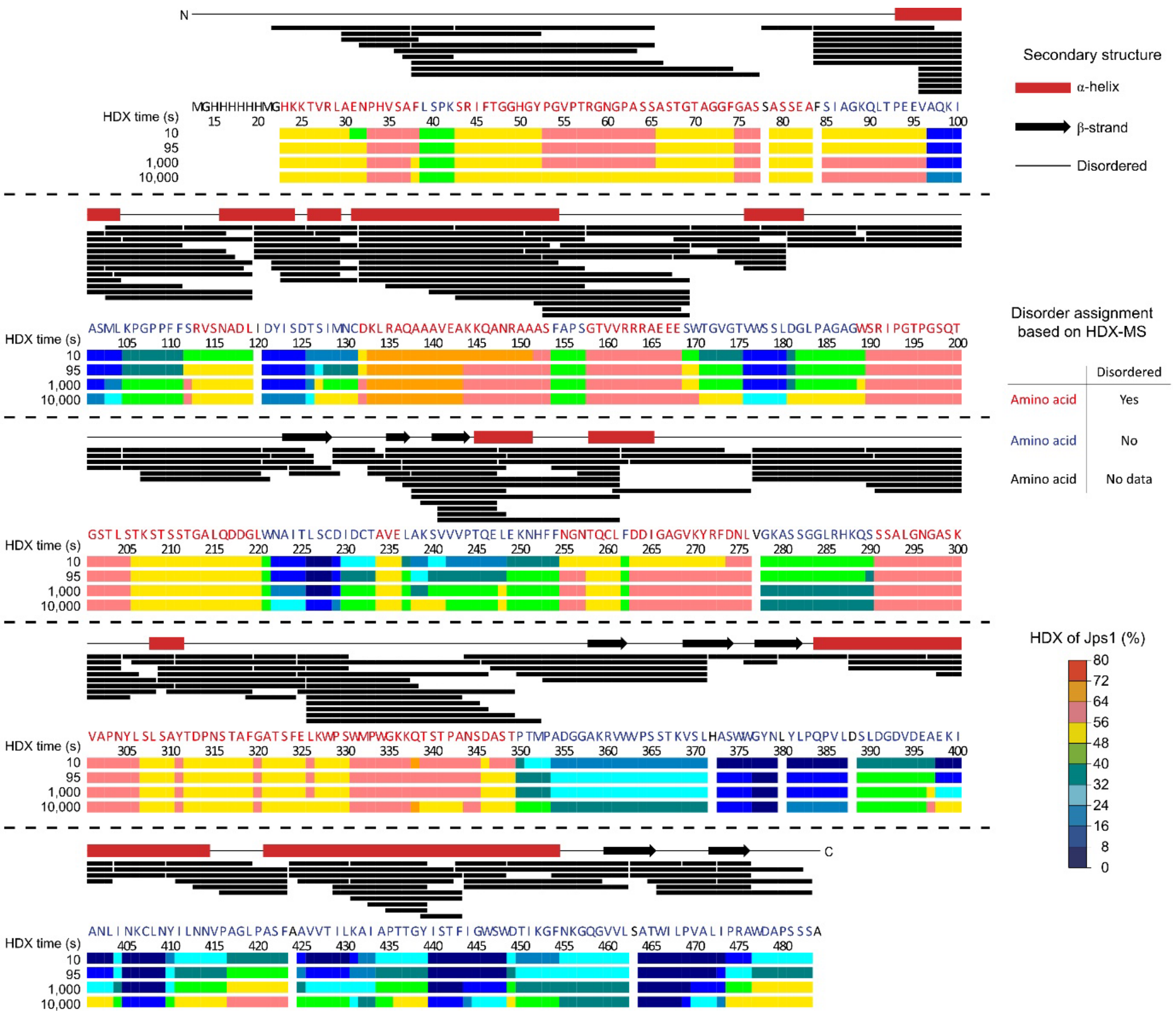

**Supplementary Figure 4. Disordered regions of Jps1 in solution probed by HDX-MS.** The residue-specific HDX of Jps1 is plotted on its amino acid sequence. Each black bar represents a peptide identified in HDX. The secondary structure as per the AlphaFold2 model of Jps1 (19) is illustrated above. Residues are colored in red or blue for residues predicted as disordered or possessing higher-order structure respectively, and residues not covered by peptides are colored black.

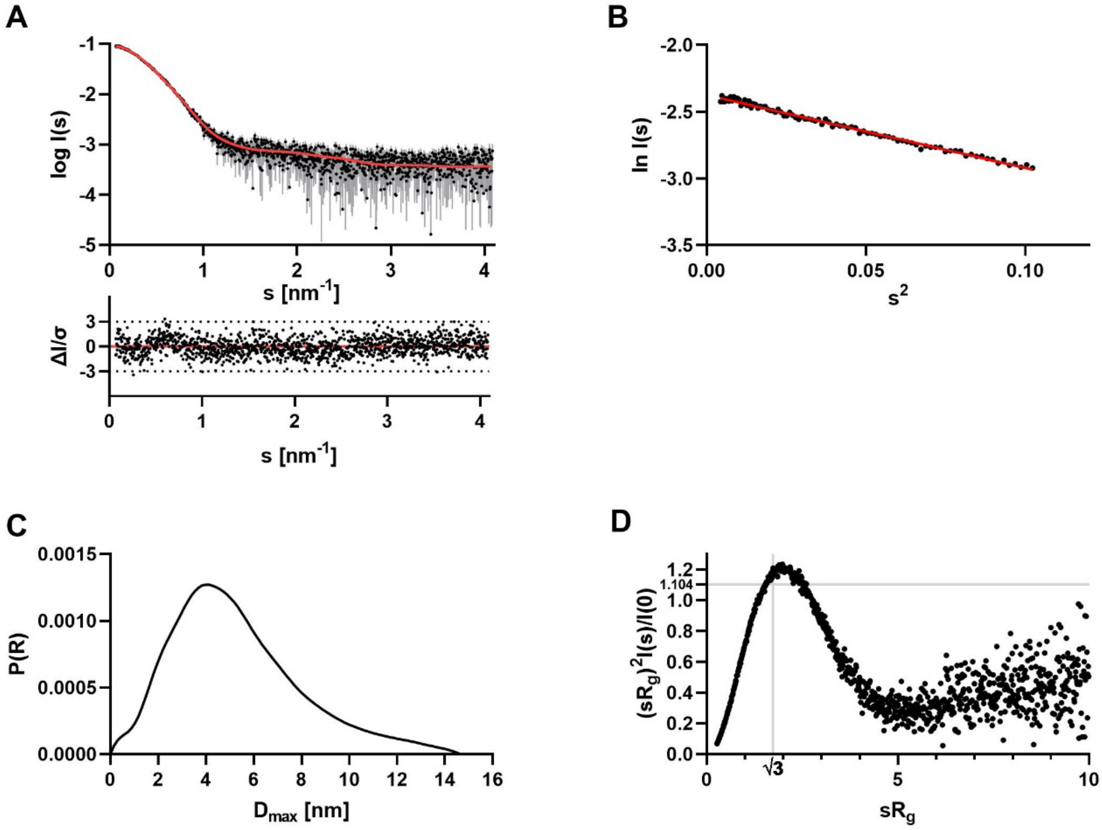

**Supplementary Figure 5. Small-angle X-ray scattering data from Jps1$^{1\text{-}484}$. A.** Scattering data of Jps1$^{1\text{-}484}$. Experimental data are shown in black dots, with grey error bars. The CORAL model fit is visualized as red line and below is the residual plot of the data. **B.** The Guinier plot of Jps1$^{1\text{-}484}$ showed a stable Guinier region with a $R_g$ of 4.05 nm. **C.** The *p(r)* function of Jps1$^{1\text{-}484}$ shows a maximum intra-particle distance ($D_{max}$) of 14.65 nm. **D.** The dimensionless Kratky plot of Jps1$^{1\text{-}484}$ revealed a compact elongated molecule.

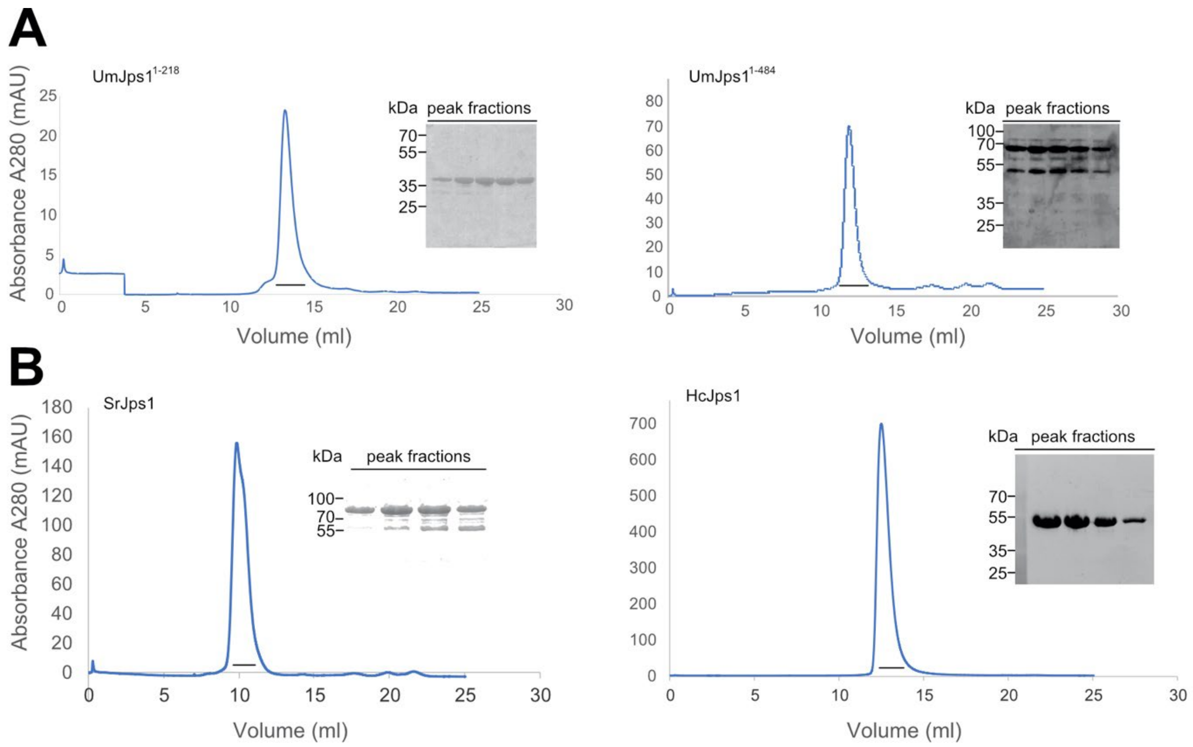

**Supplementary Figure 6. Purification of Jps1 truncations and homologs. A.** Expression of UmJps1$^{1\text{-}218}$ and UmJps1$^{1\text{-}484}$ in *E. coli*. Shown are the chromatograms obtained from size-exclusion chromatography. The Coomassie-stained SDS-PAGE show peak fractions as indicated in the chromatogram. **B.** Expression of full-length SrJps1 and HcJps1 in *E. coli*. Shown are the chromatograms obtained from size-exclusion chromatography. The Coomassie-stained SDS-PAGE show peak fractions as indicated in the chromatogram.

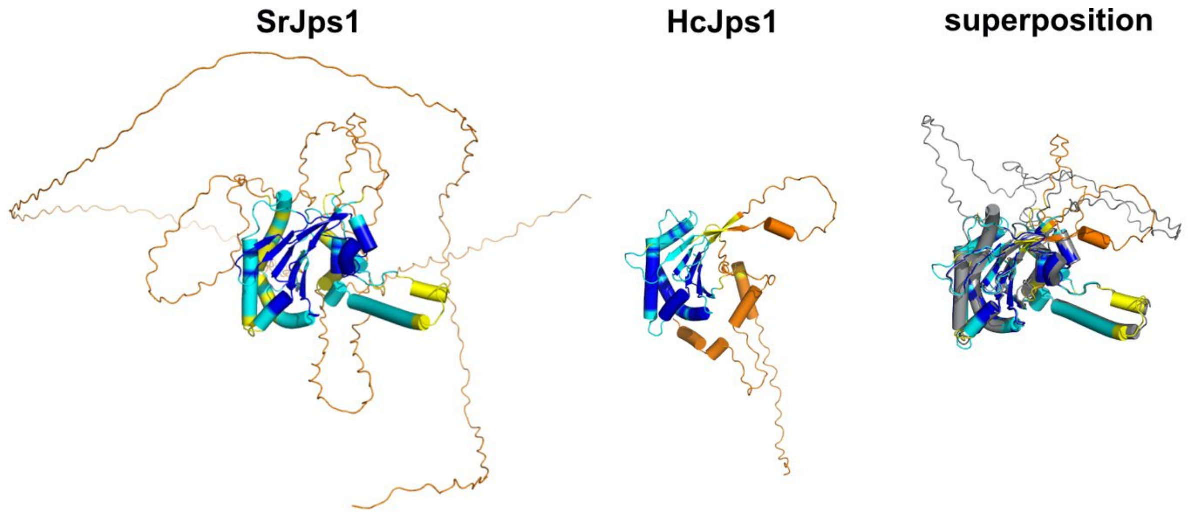

**Supplementary Figure 7. Structure prediction of SrJps1 and HcJps1.** Shown are the AlphaFold2 structural models of SrJps1 and HcJps1 colored by confidence. Structural models were superposed to UmJps1 and the flexible N- and C-termini removed to visualize that the conserved core domains align well.

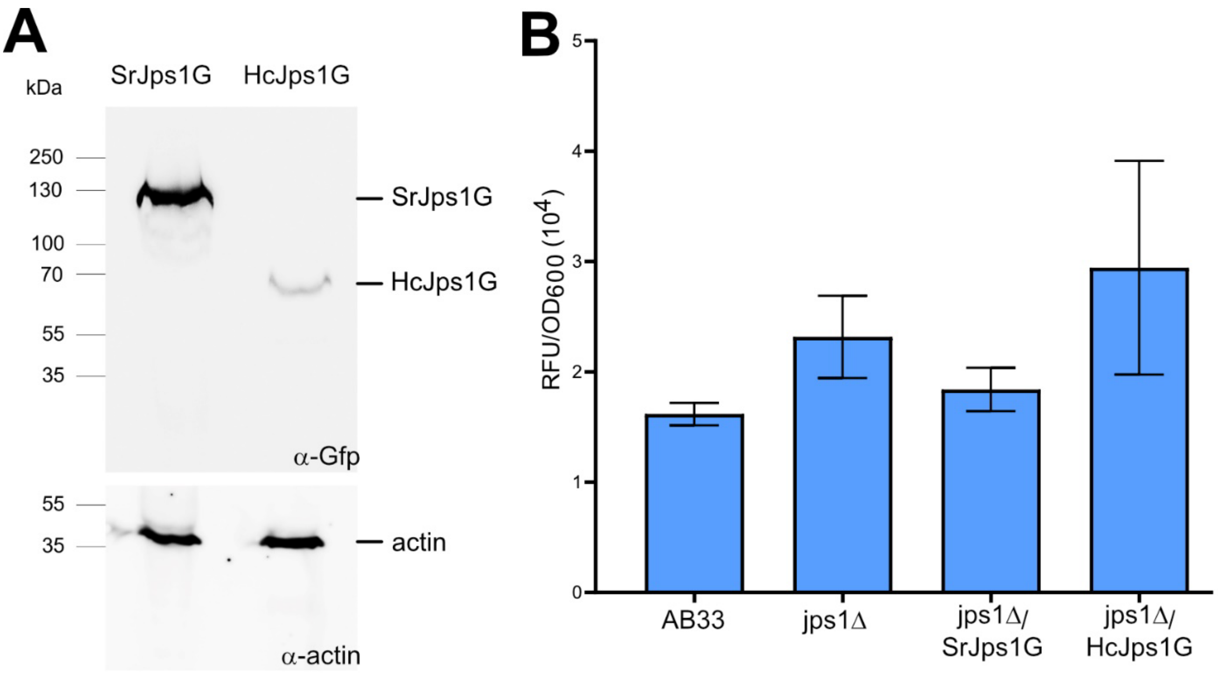

**Supplementary Figure 8. Intracellular Cts1 activity of indicated AB33 derivatives. A.** Western blots of SrJps1G and HcJps1G fusion proteins detected in cell extracts. Actin was used as loading control. **B.** Determination of intracellular chitinase activity. Background strain AB33 and AB33 jps1Δ lacking Jps1 were used as controls. Intracellular chitinase activity of AB33 jps1Δ/SrJps1G was similar to AB33 while activity of AB33 jps1Δ/HcJps1G was elevated resembling loss of Jps1. The assay was conducted in three biological replicates. Error bars indicate standard deviation.

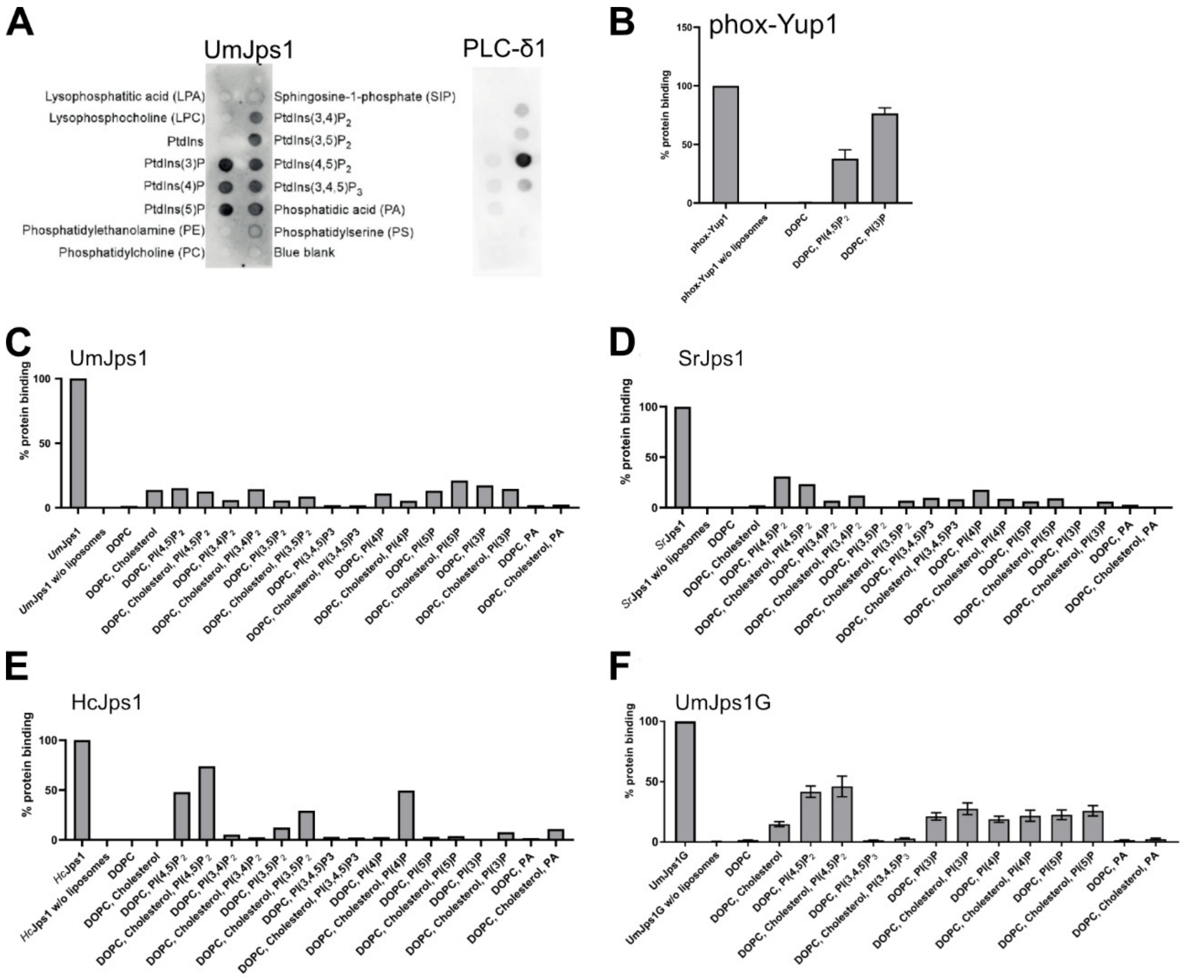

**Supplementary Figure 9. Lipid-protein interaction assays using PIP strips and liposome binding assays. A.** PIP strips were incubated with recombinant His-tagged Jps1. Antibodies directed against the His-tag were used for detection. Recombinant PLC-δ1 PH domain protein (Echelon) specifically binding PI(4,5)$P_2$ was employed as positive control. **B. - E.** Quantification of liposome binding assays using the PX domain of UmYup1 (phox-Yup1). (**B**), UmJps1 (**C**), SrJps1 (**D**), HcJps1 (**E**) and UmJps1G (**F**). Liposomes containing DOPC plus the indicated lipids were incubated with recombinant Jps1 variants. Interacting proteins localize to the pellet fraction after high-speed centrifugation. Liposome binding assays were conducted once for screening purposes (**C.-E.**), the data shown in **B** and **F** represent the mean of biological triplicates. Error bars represent standard deviation.

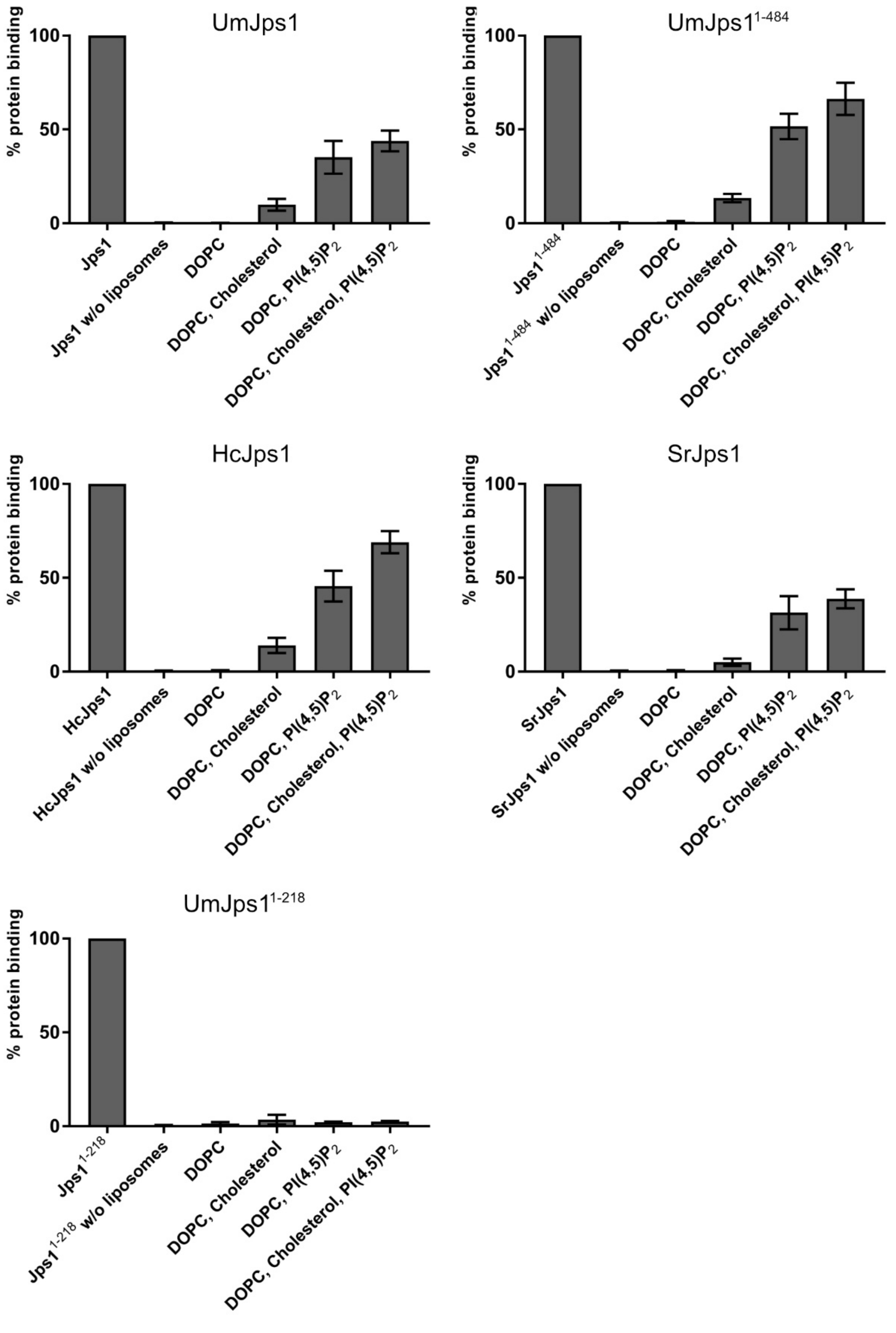

**Supplementary Figure 10. Liposome-binding assays to corroborate $PI(4,5)P_2$ interaction.** Quantification of pellet fraction from liposome binding assays using the purified proteins UmJps1, UmJps1$^{1-484}$, SrJps1, HcJps1 and UmJps1$^{1-218}$ incubated with the indicated lipids and lipid mixtures. All assays were conducted in biological triplicates. Error bars represent standard deviation.

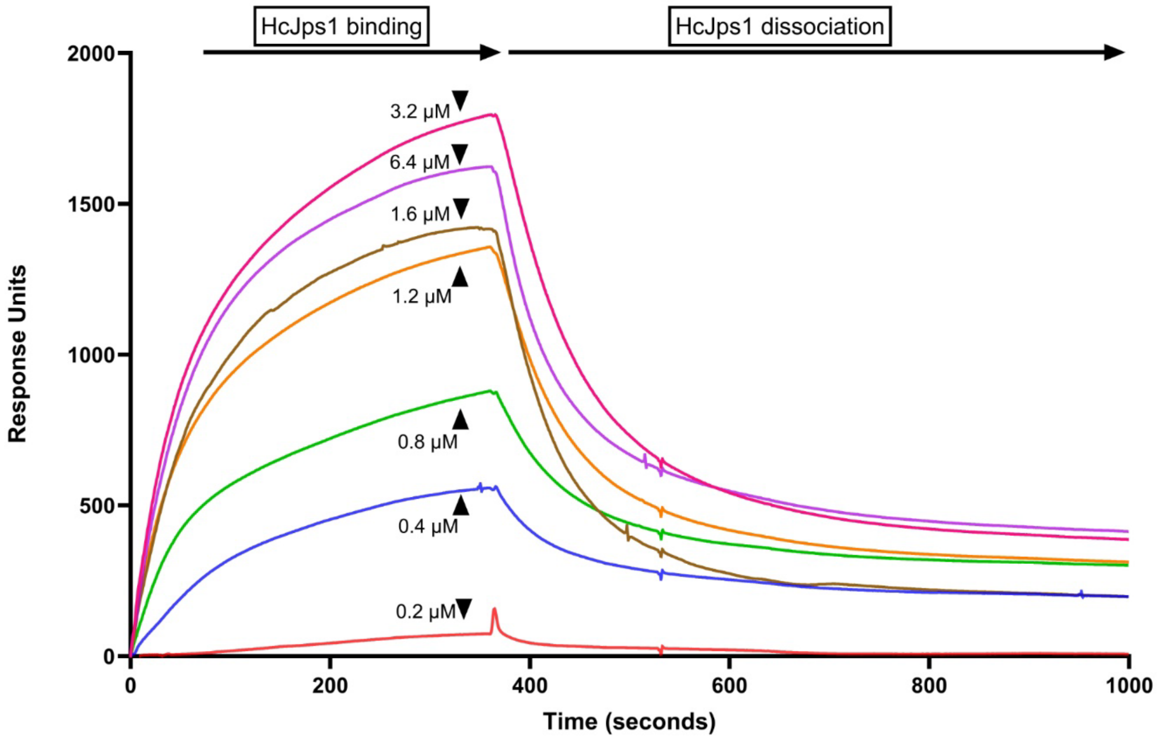

**Supplementary Figure 11. Surface Plasmon Resonance (SPR) of HcJps1.** Shown here is the concentration range of HcJps1 at 0.2 μM to 6.4 μM binding to the immobilized liposomes containing DOPC and PI(4,5)$P_2$, followed by the dissociation phase. DOPC-only liposomes were used as a negative control in the reference channel. The response units of a reference buffer (blank) was subtracted from all depicted curves.

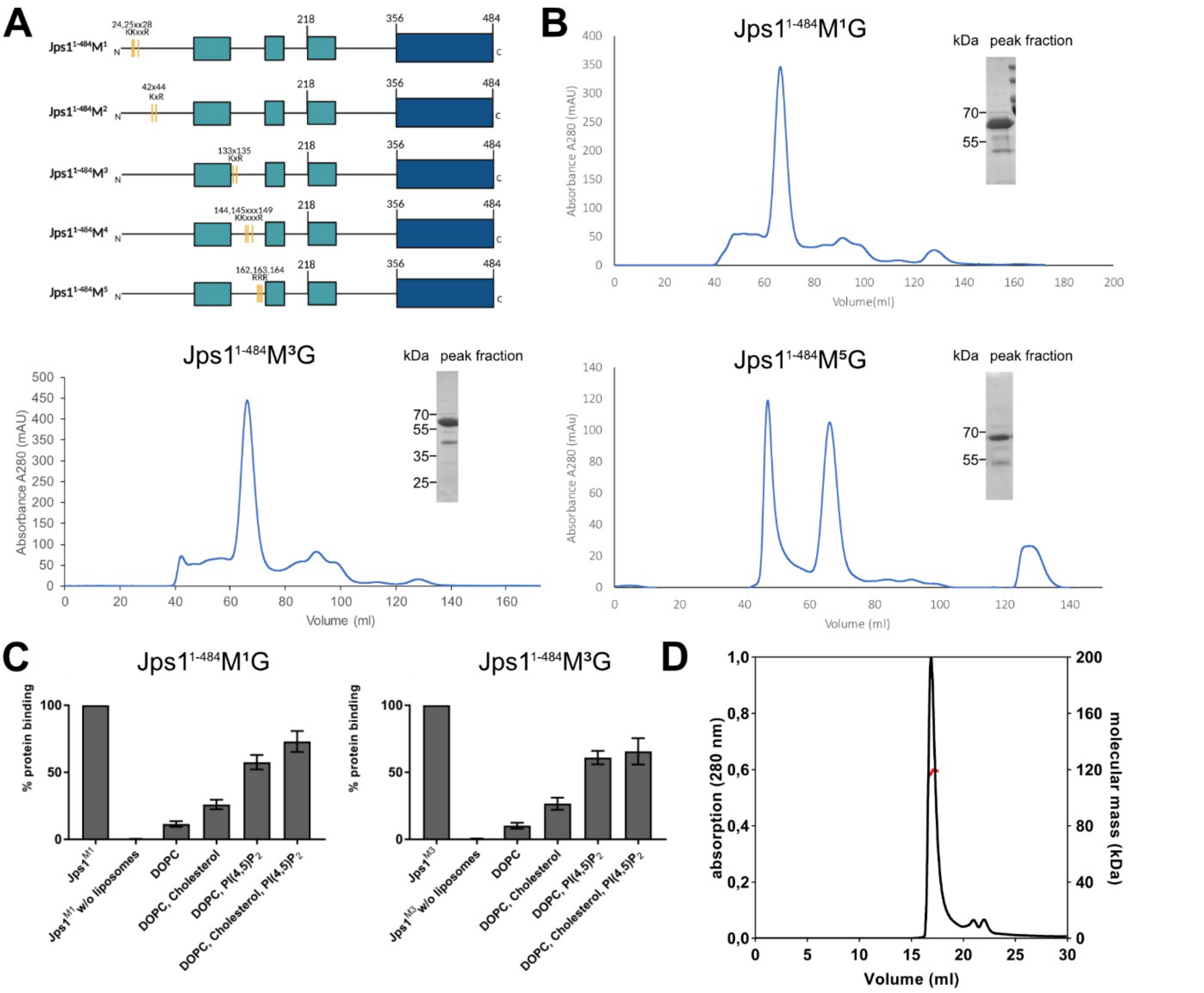

**Supplementary Figure 12. Purification and lipid-interaction assays of Jps1$^{1-484}$ variants with mutated basic clusters. A.** Schematic representation of the different Jps1 mutant versions. Mutated residues are indicated in light orange. Basic amino acids (Lysin, Arginine) were replaced by Alanine. **B.** Expression of UmJps1$^{1-484}$M$^1$, -M$^3$ and -M$^5$ in *E. coli*. Shown are the chromatograms obtained from size-exclusion chromatography. The Coomassie-stained SDS-PAGE show peak fractions as indicated in the chromatogram. **C.** Quantification of liposome binding assays. Liposomes containing DOPC plus the indicated lipids were incubated with recombinant Jps1$^{1-484}$ variants M$^1$ and M$^3$. Interacted proteins localize to the pellet fraction after high-speed centrifugation. The assay was performed in biological triplicates. Error bars depict standard deviation. **D.** SEC-MALS anaylsis of Jps1$^{1-484}$M$^5$. The black line shows the absorption at 280 nm (SEC), the molecular weight as determined by MALS is depicted in red.

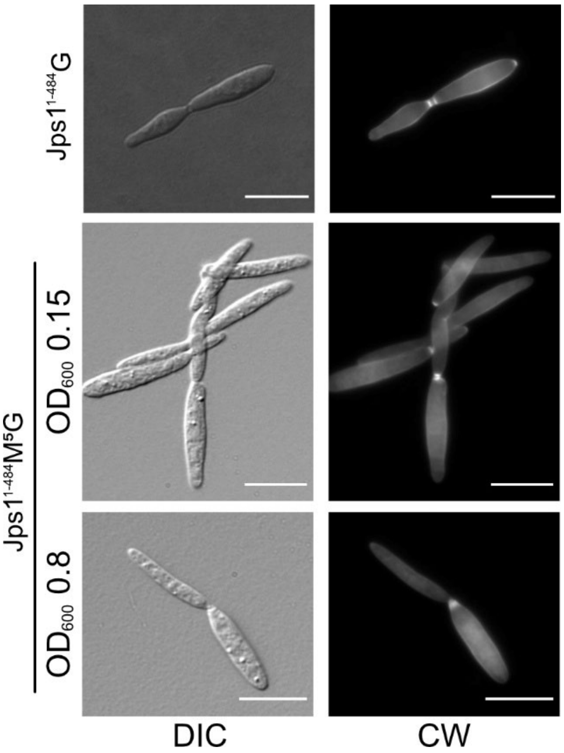

**Supplementary Figure 13. Morphology and septal staining in strains expressing $^{Jps11-484}$ and its variant M$^{5}$.** Septal staining using Calcofluor White of the AB33 jps1Δ complementation strains Jps1$^{1-484}$G, and Jps1$^{1-484}$M$^{5}$G at low and normal optical density (OD$_{600}$). DIC, differential interference contrast. CW, Calcofluor White. Scale bars, 10 μm.
